# Dominant CIZ1 fragments drive epigenetic instability and are expressed in early stage cancers

**DOI:** 10.1101/2023.09.22.558821

**Authors:** Gabrielle L. Turvey, Ernesto López de Alba, Emma Stewart, Lewis Byrom, Heather Cook, Sajad Sofi, Ahmad Alalti, Justin F-X Ainscough, Andrew Mason, Alfred A Antson, Dawn Coverley

**Author notes:** **Corresponding authors:** Gabrielle Turvey Dawn Coverley.

## Abstract

CIZ1 is a nuclear matrix protein that is part of the large RNA-dependent supramolecular assembly complexes (SMACs) that form at the inactive X chromosome (Xi) in female cells, and smaller assemblies throughout the nucleus in males and females. It plays a role in maintenance of epigenetic state and gene expression in differentiated cells, via stabilisation of histone post-translational modifications H2AK119ub1 and H3K27me3, added by polycomb repressive complexes (PRC) 1 and 2.

Here, we show that expression of the N-terminal replication domain (RD) and C-terminal anchor domain (AD) of human CIZ1 transcript is uncoupled, with consistently elevated AD in early stage breast cancers, and sporadically elevated AD in other common solid tumours. At the protein level CIZ1-Xi SMACs are corrupted in female breast cancers cells, and this is accompanied by elevated AD-encoding transcripts.

We modelled the effect of AD fragments in primary murine embryonic fibroblasts and observed dominant-negative interference with CIZ1 SMACs during their assembly in early G1 phase. Mutagenesis identified the matrin 3 homology domain as essential for self-interaction to form stable homodimers *in vitro*, and as a determinant of its dominant-negative effect in cells, implicating the dimerization interface in CIZ1 SMAC integrity.

SMAC disruption was coincident with depletion of PRC1-dependent H2AK119ub1 from Xi chromatin, in a manner abrogated by the PR-deubiquitinase inhibitor PR619, suggesting that CIZ1 SMACs normally stabilise H2AK119ub1 by shielding Xi chromatin from attack by deubiquitinases. Moreover, SMAC disruption was accompanied by changes in gene expression within days.

Together, the data suggest that inappropriate expression of CIZ1 AD fragments could drive epigenetic instability in early stage breast cancers by destabilizing the CIZ1 SMACs that normally protect repressed chromatin.

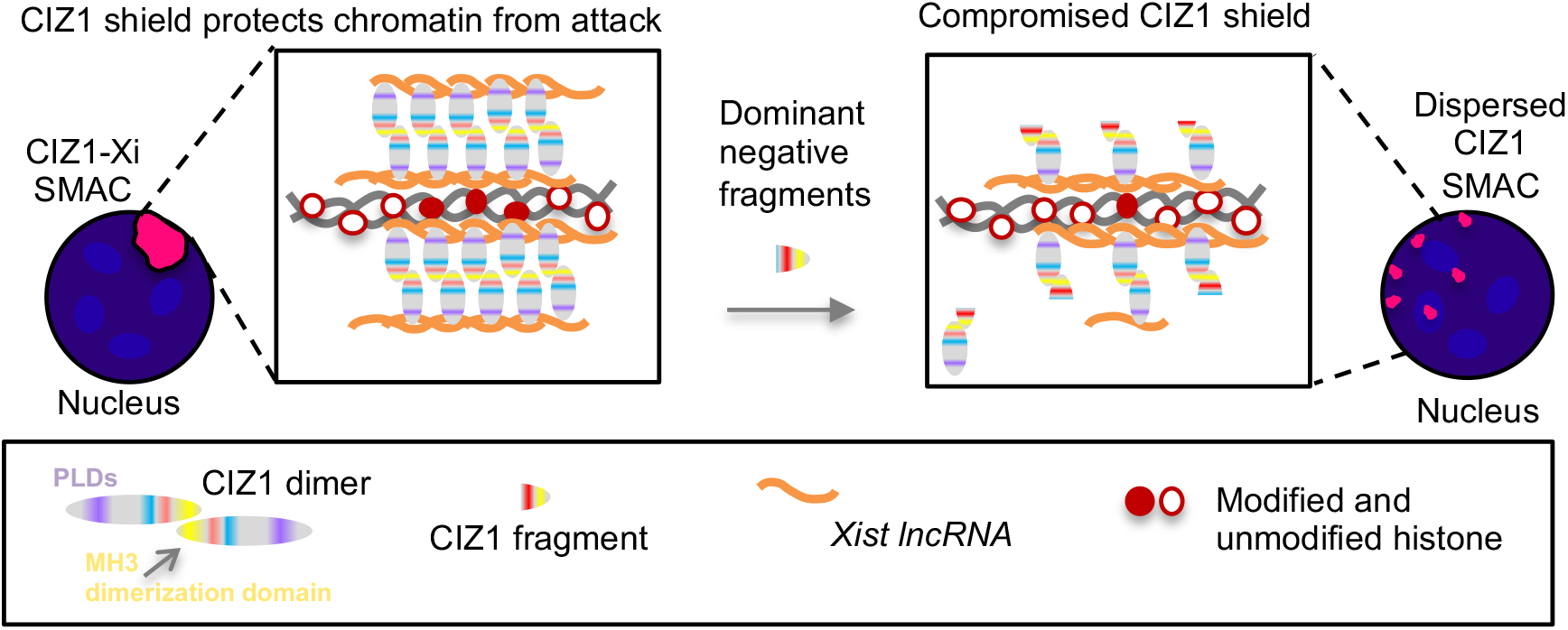

---

The process by which chromatin is selected and packaged into a transcriptionally repressed state that is stably maintained through cell generations is central to development and disease. Failure to maintain tight control creates a permissive state in which cellular responses to intrinsic or extrinsic cues is compromised, allowing inaccurate cell fates. In fact, weakened repression of heterochromatin can result in pro-oncogenic changes and has the potential to give rise to all the classic hallmarks of cancer, even in the absence of genetic change ^1, 2^. The inactive X chromosome (Xi) is the most intensely studied model of chromatin repression and is revealing how the *cis*-acting lncRNA *Xist*^3, 4^ directs the formation of large RNA-dependent supramolecular assembly complexes (SMACs) populated by chromatin modifying enzymes^5^. Aggregation of SMAC proteins, mediated by their intrinsically disordered regions (IDRs), creates a functional nuclear compartment that concentrates regulatory factors to establish local gene silencing early in development.

Cip1-interacting zinc finger protein 1 (CIZ1) is one of several proteins that populate Xi SMACs, recruited via its interaction with the repeat E element of *Xist*^6, 7^. However, several observations set CIZ1 apart from other SMAC components. First, it is not required for *Xist* recruitment, Xi silencing or embryonic development, and the impact of its loss only becomes apparent in somatic cells in which repressed chromatin is already established, but must be faithfully maintained. In differentiated fibroblasts from CIZ1 null mice, local retention of *Xist* around Xi chromatin is compromised^6, 7^, repressive histone post-translational modifications (PTMs) are lost, and genome-wide changes in the expression of genes under the regulation of polycomb are apparent^8^. Second, the stability of CIZ1 within Xi SMACs, even those that form during the initiation stages of X-inactivation, is unusually high. Compared to other SMAC components (SPEN, PCGF5, CELF1, PTBP1), the residency time of CIZ1 is estimated to be 2-10 fold longer, similar to that of *Xist*^5^. Thus, it appears that CIZ1 exchanges less readily than other protein components, and might therefore contribute a stabilizing influence on *Xist* and Xi SMACs.

Some of the sequence determinants required for formation of CIZ1-Xi SMACs are known, including two alternatively-spliced, low-complexity prion-like domains (PLD1 and PLD2) that modulate interaction with *Xist,* and a second RNA interaction domain in the C-terminus ^9^. Neither RNA interaction is sufficient alone to support incorporation of CIZ1 into Xi SMACs, but when present in the same molecule they support accumulation of CIZ1 at Xi, and drive *de novo* enrichment of H2AK119ub1 and H3K27me3, added by polycomb repressive complexes 1 and 2 (PRC1/2). These experiments indicate bi-valent interaction with RNA and show that CIZ1 SMAC formation is directly linked with modification of chromatin^9^.

Disappearance of the Barr body (Xi) has been known for decades and is considered a hallmark of cancer. Erosion of the Xi in breast tumours and cell lines was originally ascribed to genetic instability, though more recent analysis also implicates epigenetic instability, evident as abnormal sub-nuclear organization, aberrant promoter DNA methylation, and perturbations of chromatin including H3K27me3^10^. Transcriptional reactivation of X-linked genes has been implicated in both breast and ovarian cancers^11^ and may be indicative of wider, and possibly earlier, epigenetic erosion. In fact, widespread erosion of the DNA methylation landscape can give rise to the transcriptional changes common in tumors^12^, and for breast cancers in particular the progression from progenitor cell to pre-malignant lesion has been shown to involve changes in the DNA methylome that precede genetic instability^13,14^. From data such as these, a model is emerging in which induction of breast cancer could occur primarily through epigenetic disruption.

Here, we explored the aberrant expression of CIZ1 in human cancers, and find consistent over-expression of part of the CIZ1 gene in primary breast tumours, even at stage 1. We also uncovered an underpinning mechanism by which dominant-negative CIZ1 protein fragments (DNF’s), disrupt the structure and function of CIZ1 assemblies in Xi SMACs, causing loss of repressive modifications in underlying chromatin, and changes in gene expression. We suggest that disease-associated CIZ1 DNF’s contribute to epigenetic instability in human cancers by destabilizing CIZ1 assemblies, and that this may play an early role in tumour aetiology.

## Results

### CIZ1 SMACs are disrupted in breast cancer cells

In primary epithelial cells derived from normal human female mammary tissue (HMECs) a single large CIZ1 assembly is visible in approximately 80% of cells in a cycling population (Fig.1A). This is coincident with local enrichment of H2AK119ub1 identifying the assembly as at the Xi, and is dependent on multivalent interaction with RNAs including *Xist* (Fig.1B ^9^). In normal cells CIZ1 SMACs occur with similar frequency regardless of whether epitopes in its N-terminal DNA replication domain (RD ^15^), or C-terminal nuclear matrix anchor domain (AD ^16^) is detected (Fig.1C). Enrichment of CIZ1 within Xi SMACs has been shown previously in human ^6, 17^ and murine cells ^6, 7^.

**Figure 1.**
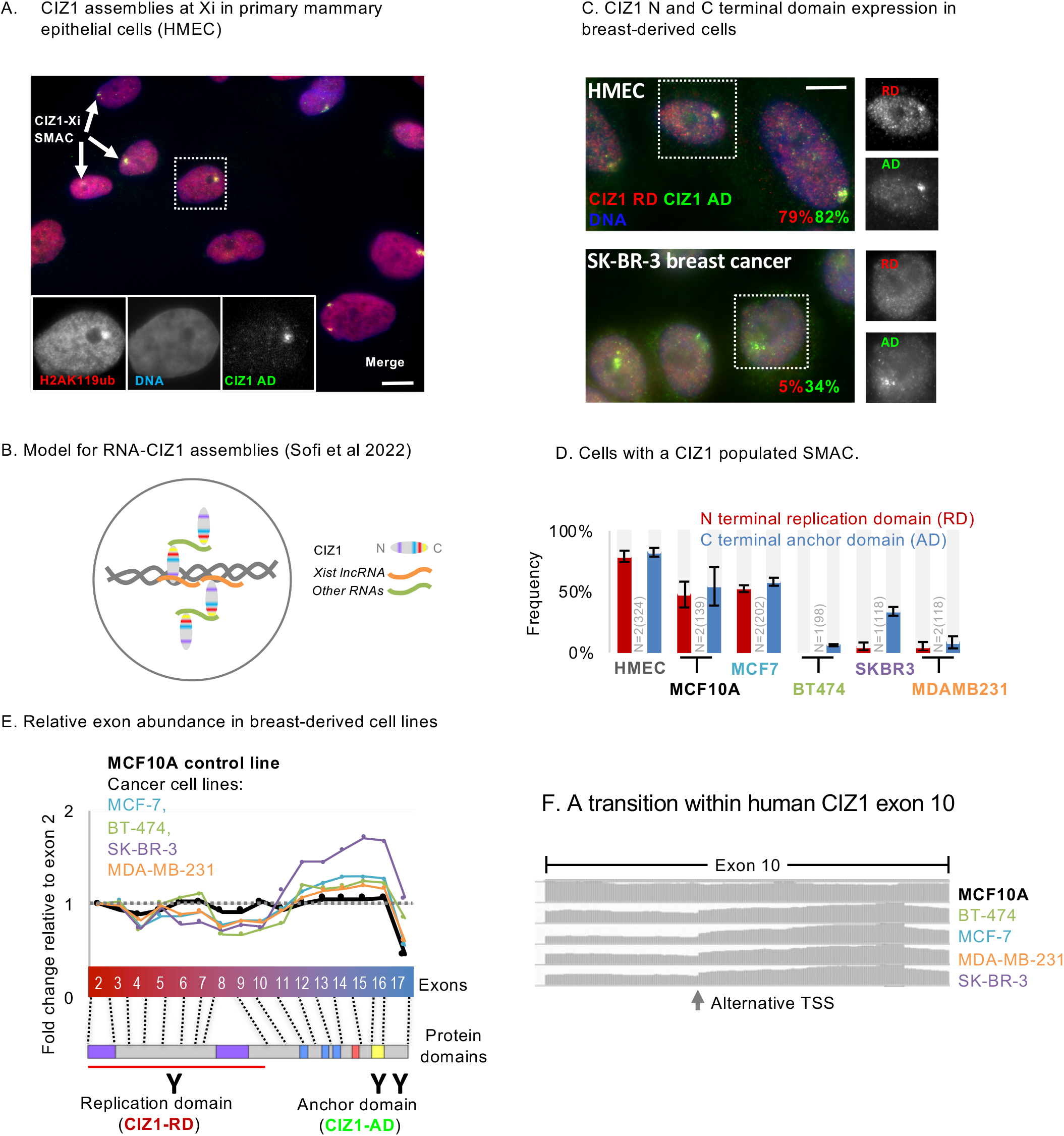
Uncoupled and imbalanced CIZ1 domain expression in breast cancer cell lines. A) Female primary human breast epithelial cells (HMECs) stained for CIZ1 via its C-terminal anchor domain (AD, green), and co-localization with H2AK119ub1 (red) as a marker of Xi chromatin. DNA is blue. Inset, example nucleus with CIZ1-AD and H2AK119ub1 shown individually in grey scale. Bar is 10μm. B) Recent model showing multivalent interaction between CIZ1 N- and C-terminal RNA interaction domains, and RNAs including *Xist* in the vicinity of the inactive X chromosome ^9^. C) Representative immunofluorescence images of CIZ1-RD (red) and CIZ1-AD (green) in HMEC and the breast cancer cell line SK-BR-3, after pre-fixation wash with detergent-containing buffer. Right, nuclei in which RD and AD are shown individually in grey scale, illustrating differential detection in the breast cancer line. Bar is 10μm. Percentage values indicate frequency of cells with CIZ1 RD and AD assemblies. Additional cell lines are in SFig.1A,B, also showing pre-fixation high-salt exposure. C) Frequency of cells with discrete nuclear CIZ1 assemblies, detected via CIZ1-AD (green, Ab87) or CIZ1 replication domain (RD, red, Ab1793) in cycling populations of the indicated breast-derived cell types. Error bars show SEM. A reduced frequency of CIZ1 SMACs is observed in non-cancer breast cell line MCF-10A and cancer cell line MCF7, while in the more aggressive BT-474 and MDA-MB-231 cancer cells, large CIZ1 SMACs are rare for both RD and AD epitopes, and in SK-BR-3 populations only detectable via the AD epitope. In all four of the cancer lines the appearance of those assemblies that are detected is less compact and coherent (see SFig.1). E) CIZ1 exon-specific TPMs from four breast cancer (MCF7, BT-474, SK-BR-3 and MDA-MB-231) and one normal breast-tissue derived cell model (MCF10A), normalised to the first translated exon (exon 2), showing imbalanced domain expression favouring anchor domain. Below, exon map aligned with protein domains (see SFig.4A), and location of epitopes detected throughout. F) Frequency of reads aligning to human CIZ1 exon 10, demonstrating consistent coverage in the normal MCF10A line, and a transition in the cancer cell lines within exon 10, at the location of an alternative transcription start site.

Compared to HMECs, in four breast cancer cell lines CIZ1 Xi SMACs are either absent, less compact and coherent, or RD and AD epitopes are differentially susceptible to extraction from the nucleus (Fig.1C,D and SFig.1A,B). This suggests that CIZ1 RD and AD are not always part of the same polypeptide. For a description of CIZ1 protein in all six breast-derived cell lines see legend to SFig.1. We conclude that CIZ1-Xi SMACs are commonly disrupted in breast cancer cell lines, consistent with the reported wider destabilization of the inactive X chromosome in breast cancer cells and tissues ^10^.

### Uneven CIZ1 expression in breast cancer cell lines

Alignment of transcriptomes from the four breast cancer-derived cell lines, and control cell line MCF10A, to CIZ1 translated exons (2-17) revealed over-representation of AD-encoding exons compared to RD-encoding exons for the tumour-derived lines (Fig.1E). We also noted a transition in transcript coverage within exon 10 (Fig.1F). This coincides with one of five predicted internal transcription start sites (TSS) annotated in Ensembl ^18^ and identified in the FANTOM5 project ^19^, and with enrichment of indicators of active chromatin (H3K4me1, H3K27ac, H3K79me2 and DNase hypersensitivity) in the cancer cell lines but not in normal HMECs (SFig.1C). Thus, archive data suggests that transcription can begin from an internal site in the CIZ1 gene, and that its use in breast cancer cells may lead to over-representation of C-terminal sequences.

To ask whether this could translate to an excess of anchor domain polypeptide, we profiled CIZ1 protein expression across the same set of cell lines by western blot (SFig.1D). CIZ1 RD antibody detected two species at 110kDa and 130kDa, as seen previously in both human and mouse derived cells, but with varying proportions, while CIZ1 AD antibody revealed a family of lower molecular weight species with greatest complexity in the breast tumour-derived lines. Taken together the data support the idea that CIZ1 RD and AD domain expression is uncoupled, and that AD is over-represented at both the transcript and protein level in human breast-derived cell lines, but also highlights considerable complexity in the family of forms produced.

### Consistent elevation of CIZ1 AD-encoding transcript in early-stage breast tumours

To measure CIZ1 transcript expression in a range of primary common solid tumours, we used quantitative RT-PCR to compare the 5’ end to the 3’ end (which contribute coding sequence to RD and AD respectively), by detection of amplicons unaffected by alternative splicing (Fig.2A). In cDNAs from 46 tissue samples (SFig.2A), correlation between two RD amplicons (exons 5 and 7), or between two AD amplicons (exons 14 and 16), was strong (r=0.89, 0.87 respectively). However, RD and AD do not correlate with each other (r=0.02, 0.17). This confirms that expression of RD and AD are commonly uncoupled, and shows that the differential can be sampled across exons 5-7 verses 14-16.

**Figure 2.**
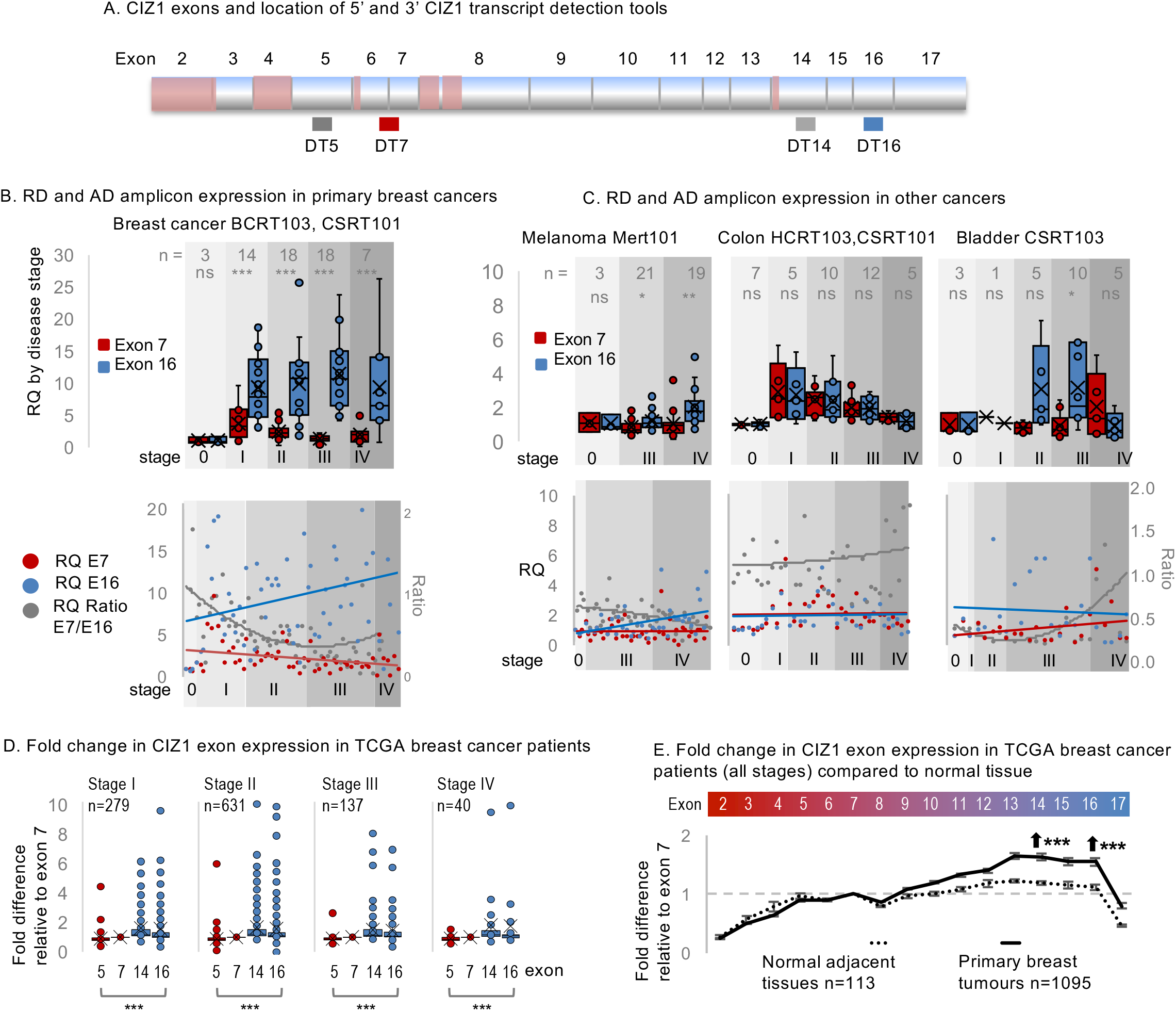
Elevated CIZ1 anchor domain expression in primary cancers. A) Exon structure of CIZ1 showing all 16 translated exons (2-17), those subject to alternative splicing (pink) and the location of amplicons detected by quantitative RT-PCR. See SFig.2A for details. B) Relative quantification (RQ) of CIZ1 exon 7 (red) and CIZ1 exon 16 (blue) in primary human breast tissue-derived cDNAs in Origene arrays BCRT103 and CSRT101 (n=60). Box and whisker plots (upper) show results aggregated by clinical stage (0-IV), where 0 represents histologically normal tissue. Dot plot (lower) shows individual samples, and the ratio between exon 7 and exon 16, with trendline (grey). Results are calibrated to the average of the stage 0 samples for each amplicon. N denotes number of samples in each class. Significance indicators show comparisons between amplicons by t-test, where ns is not significant, *p<0.05, **p<0.01, ***p<0.001. Individual sample values are given in Supplemental Dataset 1. C) As in B for human tissue-derived cDNAs in Origene arrays MERT101 (melanoma, n=43), HCRT103, CSRT101 (colon, n=39) and CSRT103 (bladder, n=24). D) CIZ1 exon expression in 1087 TCGA breast cancer samples, separated by clinical stage and normalised internally to each individual exon 7 expression. At all stages, 5’ (red) and 3’ (blue) expression is significantly different, expressed as fold change (FC). Patients with FC values above 10 are not shown to aid visualisation but are included in statistical analysis. Comparison between exon 5 and 16 is by Mann-Whitney U test, where Stage I p=0; stage II p=0; stage III p=0; stage IV p=0.000002. E) Average of all 1095 TCGA breast cancer to normal adjacent tissue-derived CIZ1 transcripts, mapped to exon 2-17 and normalized to exon 7. Breast cancer samples display elevation from around exon 10. Comparison of transcript levels in exon 5, and 14 or 16 (arrows) between normal adjacent tissues and cancer samples is by Mann-Whitney U test, where exon 14 and 16 are elevated (p=0.000004 and p=0.000175 respectively). n is number of patients, and includes 8 patients not shown in Fig.1E due to unknown breast cancer stage. Error bars show SEM.

Domain disparity was striking and consistent in breast tumours across all stages (Fig.2B). It was also evident in bladder cancer at stage III and melanoma at stage IV (Fig.2C), and sporadically in other tumours of different aetiology (SFig.2B). In addition, in some colon, lung and thyroid tumours both RD and AD domains of CIZ1 were elevated compared to histologically normal tissue (Fig.2C,SFig.2B, Supplemental Data set 1).

Focussing on breast cancer, we analysed CIZ1 expression in 1095 transcriptomes submitted to The Cancer Genome Atlas (TCGA). While transcripts that map to the whole CIZ1 gene (ENSG00000148337) revealed no *overall* difference in expression between tumours and normal tissue (significance threshold log_2_ fold change >1, p<0.05, SFig.3A), the same raw data when mapped to individual CIZ1 exons showed that RD (exons 5 and 7) is significantly underrepresented compared to AD (exons 14 and 16) at all stages (Fig.2D, SFig.3B). Elevation was notable from around exon 10, and significantly less marked in adjacent normal tissue than in breast tumours (Fig.2E), or the subset with patient-matched samples (SFig.3C). These data suggest that the C-terminal CIZ1 AD is over-represented in breast cancers from stage I onwards, and poses the question of whether inappropriate AD expression is functionally relevant.

### *In vitro* modelling of the effect of CIZ1 AD on SMACs

The multivalent nature of CIZ1’s interaction with RNA, and the requirement for both domains for the assembly of CIZ1 into SMACs at the Xi ^9^, lead us to hypothesize that fragments of CIZ1 encoding only one of its RNA interaction interfaces might have a destabilizing effect. Moreover, based on what we know of CIZ1 genetic deletion and the co-dependency of CIZ1 and *Xist* ^5–7, 20^, we hypothesized that interference with CIZ1 SMAC integrity would affect Xi chromatin. We initially modelled this in murine cells after ectopic expression of the C-terminal 275 amino-acids of murine CIZ1 (from exon 10 onwards), here referred to as C275 (Fig.2A). Human and murine CIZ1 possess the same conserved domains encoded by the same exons in the same order, and are 65% identical at the protein level, with identity concentrated in the conserved domains (up to 96%, SFig.4). Thus, we expect that functional insight uncovered in the murine system will be relevant in human cells.

In short-term (one cell cycle) transient transfection experiments endogenous CIZ1 Xi SMACs were diminished or lost in cells expressing GFP-C275 (Fig.3B) but, as seen previously ^9^ GFP-C275 did not itself accumulate at Xi. In untransfected cells (UT) in the same population, or parallel populations expressing empty GFP vector (EV) CIZ1 SMACs were unaffected. This was evidenced via detection of CIZ1 RD epitope (which is not in C275) in multiple independent cycling cell populations, and is expressed as SMAC frequency. Deletion of two zinc fingers to produce the smaller C181 fragment, did not diminish the disruptive effect on endogenous CIZ1 SMACs (Fig.3B). Thus, C-terminal fragments of CIZ1 do indeed have the capacity to interfere with endogenous CIZ1 Xi-SMACs in female cells, and will be referred to as CIZ1 DNF’s (dominant-negative fragments).

**Figure 3.**
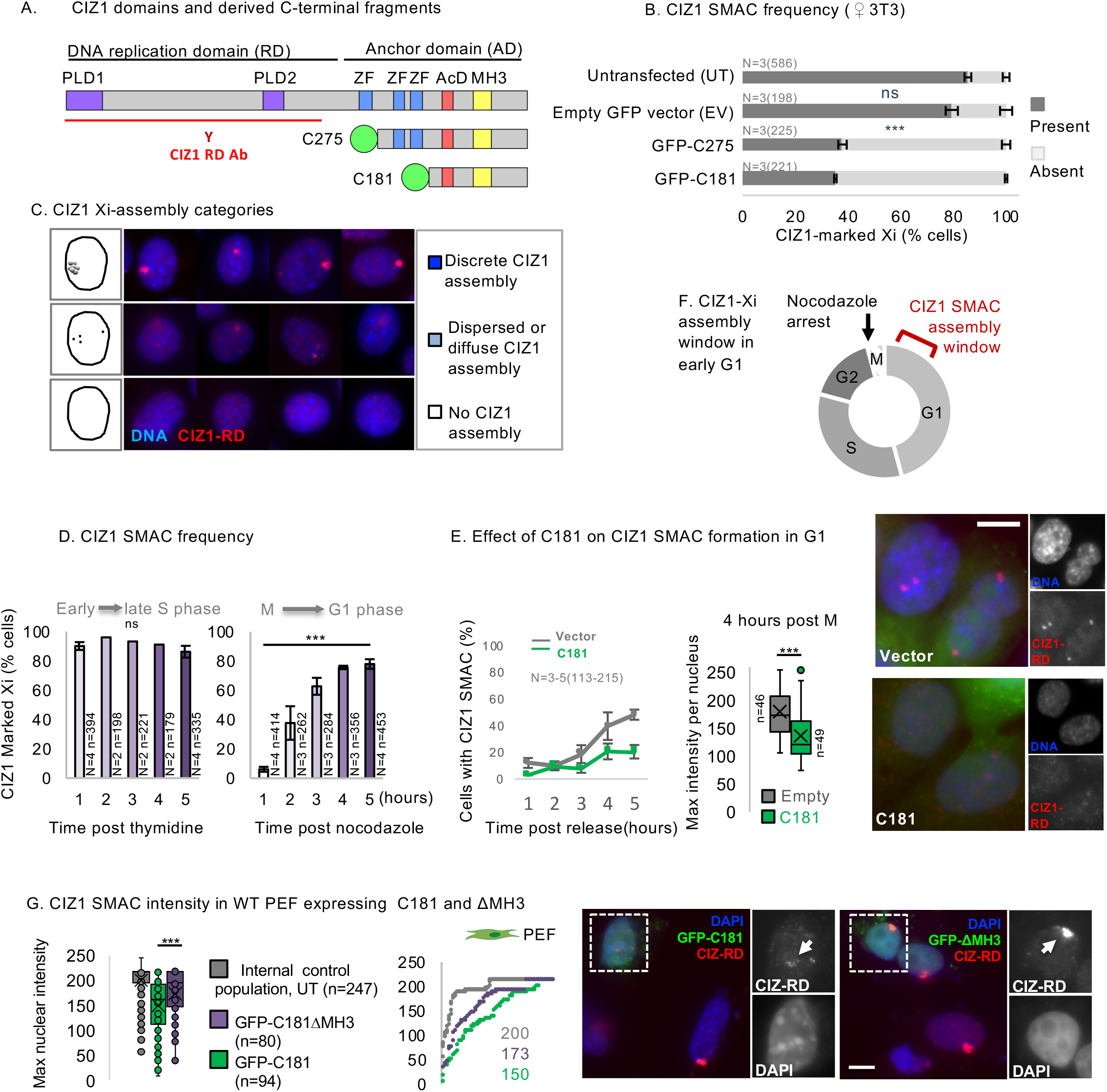
Dispersal of endogenous CIZ1 Xi SMACs by ectopic CIZ1 fragments. A) Diagram of murine CIZ1 full-length protein (ref. seq. NP_082688.1) showing prion like domains (PLD) 1 and 2 ^9^, three zinc fingers (ZF), an acidic patch enriched in aspartate and glutamate residues (AcD), and a Matrin 3 homology domain (MH3). Domain boundaries and comparison with human CIZ1 is given in SFig.4. Below, C-terminal fragments C275 and C181 correspond to the terminal 275 and 181 amino-acids respectively, both bearing an N terminal GFP tag that are used to model cellular effects. The CIZ1 RD antibody (red) was raised against a fragment of CIZ1 outside of C275 ^15^, so detects only endogenous CIZ1 in transfection experiments. B) Frequency of cells with CIZ1 Xi SMACs in untransfected (UT) populations, compared to those expressing empty GFP vector (EV), and those expressing C275 or C181. N is replicate analyses, with total nuclei inspected shown in parentheses. Comparisons are by one-way ANOVA with a Tukey post hoc test. Between EV cells and, UT cells p=0.12, C275 cells p=4.42×10^-7^, C181 cells p=2.55×10^-7^, and between C275 cells and C181 cells p=0.58. Error bars show SEM. C) Example images of female D3T3 cells taken under identical parameters, showing three categories of nuclei with either large discrete CIZ1 SMACs (upper), no detectable CIZ1 SMAC (lower), or intermediate assemblies (middle), which include those that are either dispersed into multiple smaller foci, or diminished in overall size or intensity. CIZ1 is red, DNA is blue. D) CIZ1 SMAC frequency (AD) 1-5 hours after release of female D3T3 cells from cell cycle arrest in S phase (thymidine) or M phase (nocodazole). N is 2-4 as indicated, number of nuclei inspected at each time point is given (n). Comparisons between 1 and 5 hours in each set by t-test, where p=0.45 for S phase and p=1.15×10^-6^ for M-G1 phase. E) Delayed reformation of CIZ1 SMACs after release from arrest in M phase in D3T3 cells transduced with C181 compared to vector control. Left, CIZ1 SMAC frequency 1-5 hours post release. Middle, box and whisker plot showing fluorescence intensity per nucleus at 4 hours, where n indicates the number of nuclei measured in each group. Mann-Whitney U test, p=6.1×10^-6^. Right, example images of transduced cells. Bar is 10μm. F) Illustration showing time window of CIZ1 Xi SMAC assembly early in G1 phase and the point of cell cycle arrest after exposure to nocodazole. G) Left, box and whisker plot showing endogenous CIZ1 fluorescence intensity per nucleus in female WT primary embryonic fibroblasts (45.1fc p1), untransfected (UT) or with C181, or derived deletion mutant ΔMH3, showing reduced potency of ΔMH3 compared to C181 (p=0.002, t-test). Middle, mean intensity measures ordered low to high, for endogenous CIZ1 (RD) in UT, C181 and ΔMH3 expressing cells. Mean values are also shown. Right, example images of cells stained for endogenous CIZ1 (red), with and without ectopic GFP-C181 or GFP-ΔMH3. Bar is 5μm.

Related analysis in male NIH3T3 cells using C275 also revealed a dominant-negative effect, in this case on the formation of chromatin-independent (DNase-resistant) sub-nuclear assemblies of GFP-CIZ1 (SFig.5A). This argues that the disruptive effect of CIZ1 DNFs is not limited to the super-assemblies that form at Xi, or to female cells.

### CIZ1 SMAC dispersal is cell cycle dependent

Notably, not all cells expressing CIZ1 DNF’s lose endogenous CIZ1-Xi assemblies; typically at 24 hours after transfection 30-40% of cells remain refractory to their effect (Fig.2B), and the extent of SMAC dispersal is variable. To refine quantification, CIZ1 SMACs were categorised into three distinct phenotypes; cells with a discrete SMAC, cells with no SMAC, or cells with intermediate, dispersed or diminished CIZ1 assemblies (Fig.3C), and further quantified via measurements of fluorescence density in standardised image sets (SFig.5B).

To test whether the cell cycle stage of recipient cells contributes to the heterogenous response we introduced C181 to newly contact-inhibited (arrested) cells, and observed no reduction in CIZ1 SMACs (SFig.5C). This implies a window in the cell cycle when SMACs are vulnerable to DNF’s. We also noticed that cell cycle arrest facilitates DNF accumulation at pre-existing sites of CIZ1 SMACs (SFig.5D), possibly reflecting an extended period of stability in non-cycling cells. Together the observations suggest that passage through the cell cycle offers a window in which DNF’s can exert their effect.

As in HMECs (Fig.1A), typically around 80% of murine 3T3 cells in a cycling population contain a clear CIZ1 SMAC (Fig.3B, SFig.5C). Since we know that, like *Xist* ^21^, CIZ1 SMACs disassemble in mitosis ^6^ we postulated that those cells without CIZ1 SMACs have yet to rebuild them and are in early G1 phase. We confirmed this in 3T3 cells that were synchronised in mitosis using nocodazole, and found that maximal SMAC frequency was reached by four hours after mitotic exit (Fig.3D). To look at the effect of C181 on SMACs that are forming during this window, we turned to a lentiviral transduction system to maximise delivery. When applied to cells released from mitosis, formation of SMACs was significantly delayed and those that formed had reduced CIZ1 fluorescence density (Fig.3E). This suggests that the dispersive effect of CIZ1 DNF’s is potent during the SMAC assembly window in early G1 phase (Fig.3F).

### Role of the MH3 homology domain in dimerization and SMAC dispersal

To refine the sequence requirements for SMAC dispersal by DNF’s we evaluated a set of deletion constructs, based on C181 (SFig.6A). These exclude the MH3 domain (conserved domain ZnF_U1 smart00451), or the fully human/mouse conserved sequence downstream of the MH3 domain (ΔNALTALF), or the murine equivalent of the 8 amino acids previously implicated in lung cancer (CIZ1B) ^22^, or the terminal 37 amino acids (Δ37). We also evaluated a fragment encompassing the MH3 domain but lacking sequences up and downstream (I122) ^16^. All fragments accumulated in the nucleus and were incorporated into detergent-resistant structures to differing degrees (SFig.6B). All C181 deletion mutants retained similar capability to interfere with endogenous CIZ1 SMACs, with the exception of the ΔMH3 fragment which had a small but consistent reduction in activity (SFig.6C). Diminished activity was confirmed by measuring fluorescence density in cells expressing C181 or ΔMH3 (Fig.3G). This reduced DNF activity implicates the MH3 domain in integrity of endogenous CIZ1 SMACs.

A similar set of C181 deletion mutants were expressed and purified in bacteria (SFig.6D). Size exclusion chromatography (SEC) indicated that C181 exists in a stable multimeric state (Fig.4A). Deletion of the C-terminal 37 amino acids had minimal effect on elution time, so does not contribute to multimeric state, however deletion of the MH3 domain had a significant effect. C181 was confirmed to exist as a homodimer by SEC coupled with multiple angle laser light scattering (SEC-MALLs). This generated a molecular weight estimate of approximately 40kDa, consistent with dimerization of the 20 kDa C181 monomer (Fig.4B). In contrast the ΔMH3 mutant exists as a 16kDa monomer, confirming the MH3 domain as required for dimerization. Individual full SEC and SEC-MALLs traces are given in SFig.6E,F). Consistent with this, modelling of C181 secondary structure using AlphaFold2 with instruction to form a dimeric entity ^23^, highlighted a β-strand within each MH3 domain as the putative dimerisation interface (Fig.4C). Thus, Alphafold predictions are consistent with our experimental findings, and also suggest that CIZ1 molecules are arranged in an anti-parallel orientation. Notably, the RNA interaction capability previously shown to be inherent in the C181 fragment ^9^, is diminished in the ΔMH3 mutant evidenced by reduced complex formation with *Xist* repeat E RNA probe in electrophoretic mobility shift assays (EMSA, Fig.4D), implicating dimerisation in the creation of an RNA interaction interface.

**Figure 4.**
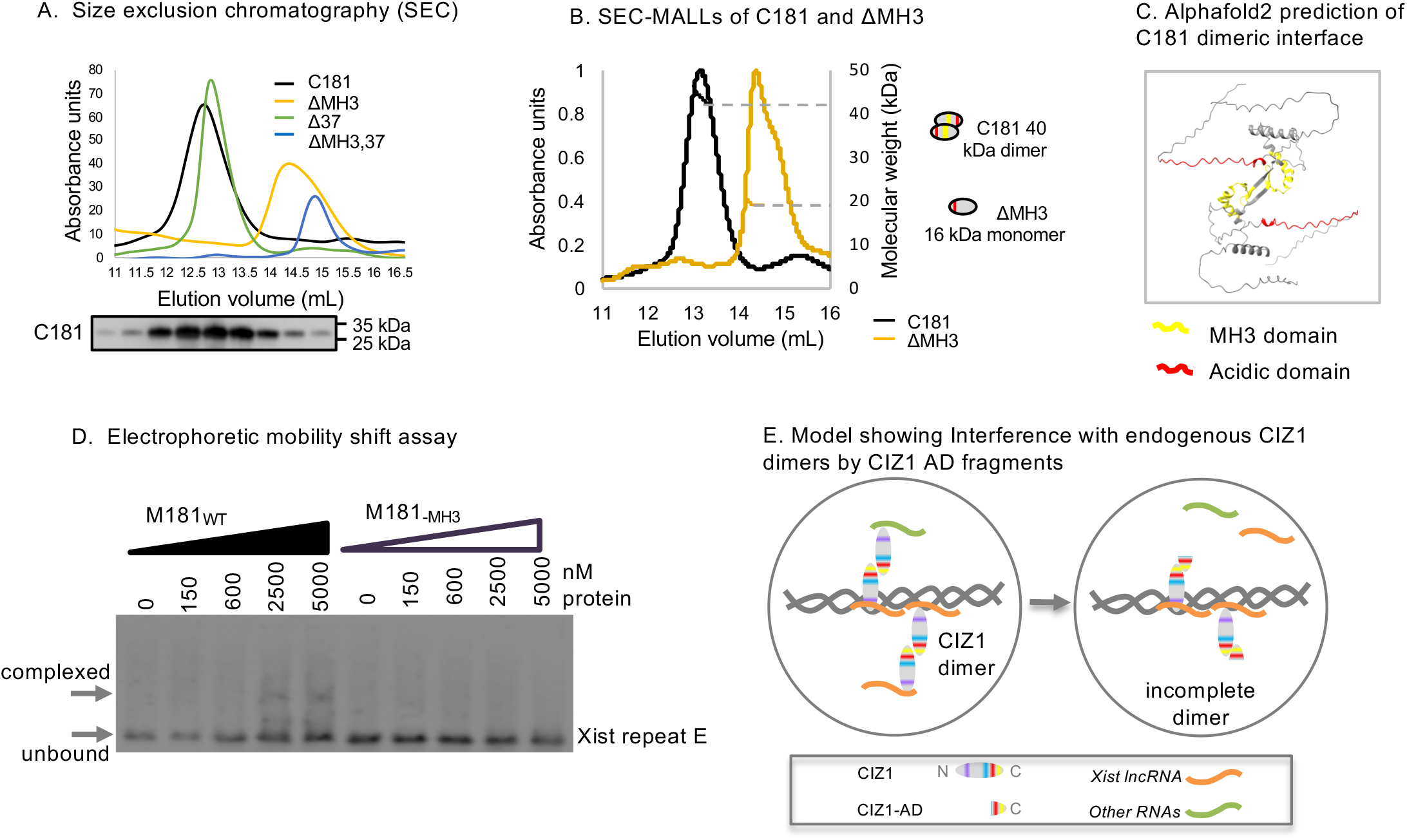
CIZ1 dimerisation via MH3 homology domain. A) Size exclusion chromatography (SEC) of the indicated recombinant CIZ1 protein fragments (see SFig.6). Compared to C181 elution time is increased for fragments lacking the MH3 domain, indicating smaller atomic radius. Below, example western blot of C181 elution fractions, probed with CIZ1 exon 17 rabbit antibody (SFig.4). B) Size exclusion chromatography multiple angle laser light scattering (SEC-MALLs) of C181 and ΔMH3, showing that C181 exists as a dimer under native conditions, whereas ΔMH3 exists as a monomer. C) Alphafold2 ^23^ modelling of C181 as a dimer, which predicts that the MH3 domain is an interface for dimerisation in anti-parallel orientation. D) Electrophoretic mobility shift assay with *Xist* repeat E RNA probe showing complex formation with C181 but not C181(ΔMH3) protein. E) Model showing interference with dimerization of CIZ1 in Xi SMACs, through the dominant-negative action of CIZ1-AD fragments.

To facilitate analysis of sequence requirements and epitope availability within CIZ1 C-terminus, we mapped a set of antibodies and determined their ability to reveal CIZ1-Xi SMACs (SFig.4B-D). These show that, *in vivo,* epitopes encoded by exon 16 (MH3 domain) are not available under standard immunofluorescence conditions, suggesting that in all CIZ1 molecules in Xi SMACs it is masked, and may therefore be in MH3-domain dimeric form. Together these results lead us to suggest that interference with endogenous CIZ1 assemblies inside cells by DNF’s may involve competition for self-interaction, and interruption of dimerisation (Fig.4E).

### Dispersal of CIZ1 SMACs leads to instability of H2AK119ub1

We postulated that dispersal of CIZ1 SMACs by DNF’s might mimic the effect on Xi chromatin seen in genetically CIZ1 null primary embryonic fibroblasts (PEFs, cultured up to passage 5). In these cells H3K27me3 and H2AK119ub1 are both depleted from Xi chromatin, and control over PRC target genes is relaxed ^8^. We also draw attention to an informative complication that arose from this analysis. Upon further rounds of cell division in culture (beyond passage 5), upregulation of enhancer of zeste homolog 2 (EZH2) the catalytic subunit of PRC2, and re-establishment of both H3K27me3 and H2AK119ub1 on Xi chromatin was observed, so that CIZ1 null cell populations resembled WT with respect to histone status at Xi. We suggested that such compensatory changes would likely be accompanied by off-target effects, and could be responsible for the observed shift towards less controlled and coherent gene expression ^8^.

To ask whether a similar chain of events (deprotection of chromatin, upregulation of modifiers, unfocussed gene expression) might be initiated by CIZ1 DNF’s, we first evaluated H3K27me3 and H2AK119ub1 in Xi chromatin in WT female 3T3 and PEFs after transient transfection with C181 (SFig.7). After 24 hours we observed no effect on H3K27me3, but a marked reduction in H2AK119ub1 in both cell types. Survival of H3K27me3 under conditions where H2AK119ub1 is depleted is consistent with analysis of the Drosophila HOX gene cluster which showed that passage through S phase is required for H3K27me3 to be diluted ^24, 25^, and with our own analysis which correlates maintenance of H3K27me3 with replication-linked relocation of Xi chromatin ^8^.

In longer term experiments, using lentiviral transduction of C181 into female WT PEF’s (Fig. 5A, SFig.7D,E), CIZ1 SMACs, H3K27me3 and H2AK119ub1 enrichment at Xi were all reduced, whether quantified by frequency, or by fluorescence intensity (Fig.5B). Together these two approaches in two cell types show that destabilization of CIZ1 Xi SMACs by DNFs, has a coincident impact on chromatin.

**Figure 5.**
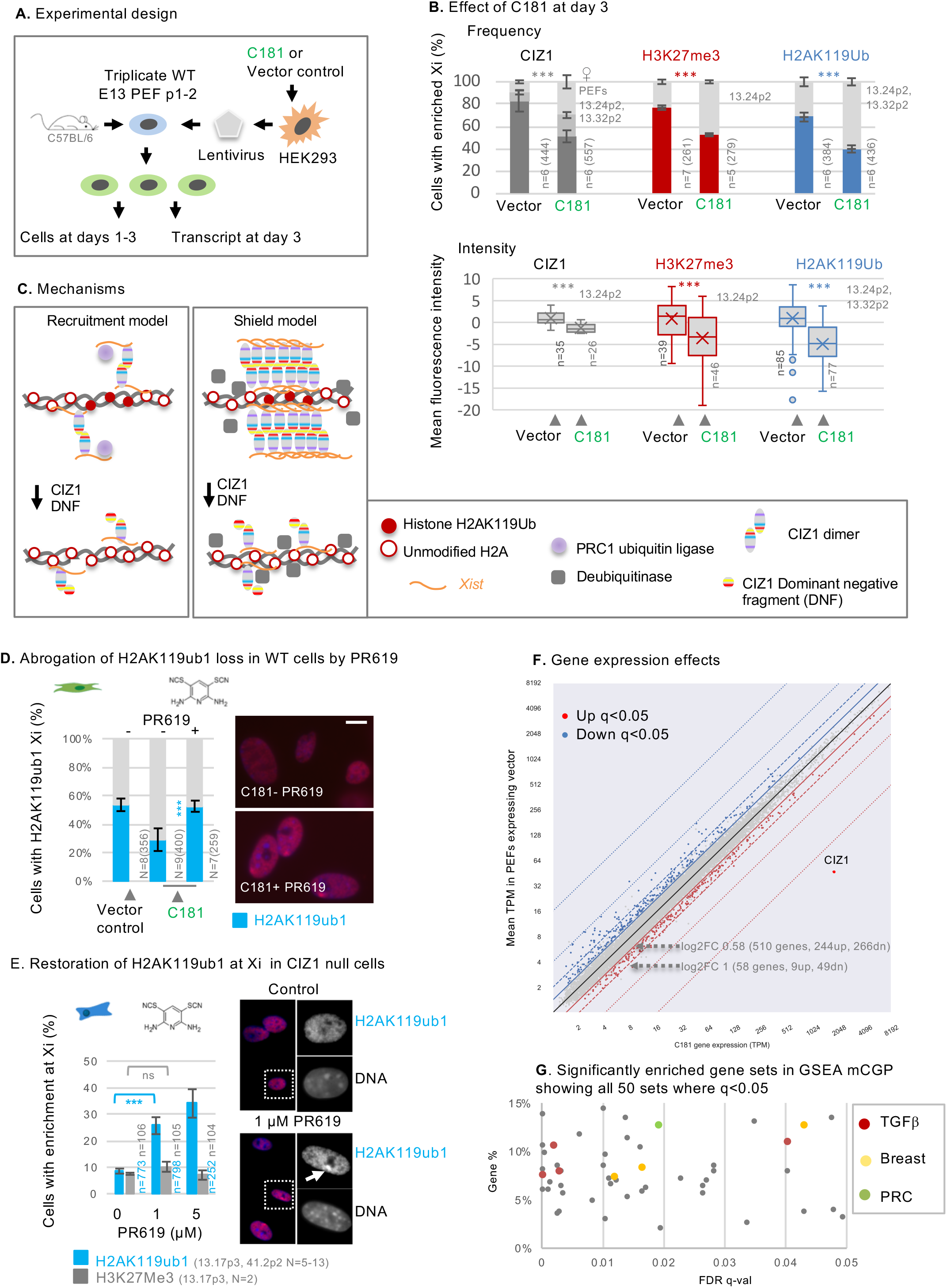
Effect of CIZ1 dominant negative fragment on histones and gene expression. A) Experimental design: polycistronic plasmids encoding ZsGreen only or ZSGreen and C181 are transfected into Lenti-X™ 293T human embryonic kidney (HEK) cells to produce lentivirus, then applied to murine primary embryonic fibroblasts (PEFs) at passage 1-2 to generate C181 or vector control transduced cells, and sampled across three days. B) Comparison of vector only populations to those transduced with C181 showing, frequency of cells with CIZ1 Xi SMACs (grey, three catagories), or H3K27me3 (red) or H2AK119ub1 (blue). PEF cell populations are listed in grey, n denotes replicate analyses with total nuclei inspected shown in parentheses. Comparisons are by a two-sample unpaired t-test, where p <0.001 in all cases. Error bars show SEM. Below, box and whisker plots showing mean nuclear intensity measures for cells transduced with C181 or vector control. C) Possible mechanisms by which CIZ1 SMACs might influence H2AK119ub1 dynamics on Xi chromatin. Recruitment model: CIZ1 dimerisation and *Xist* binding contribute to recruitment or activation of PRC1, supporting H2AK119ub1 deposition. Shield model: Multiple CIZ1 dimers and associated RNAs coalesce to form a shield at the Xi which blocks access to deubiquitinating enzymes, supporting H2AK119ub1 preservation. D) Frequency of cells with H2AK119ub1 at the Xi in the vector only population, and cells transduced with C181, without and with the DUB inhibitor PR619 (5 μM). N is replicate analyses, with total nuclei inspected in parentheses. Comparisons are by a two-sample unpaired t-test. Error bars show SEM. Right, example images taken under standardised conditions showing H2AK119ub1 in red in C181 transduced WT primary embryonic fibroblasts (strain 13.27p2). E) Restoration of H2AK119ub1 enrichment at Xi in CIZ1 null primary embryonic fibroblasts by PR619. In untreated cells approximately 10% of cells have H2AK119ub1 enriched Xi’s, which increased to approximately 35% within 24 hours of treatment, while H3K27me3 remains unchanged. N is replicate analyses, n is nuclei scored, strains and passage numbers are shown. Error bars show SEM. Right, example images of H2AK119ub1 in CIZ1 null primary embryonic fibroblasts. F) Scatter plot showing genes affected by CIZ1 DNF C181 compared to vector, based on fold change in mean transcripts per million reads (TPM), coloured blue (down) and red (up) for genes meeting the false-detection rate corrected q values of <0.05. G) Gene set enrichment analysis performed using GSEAPY on pre-ranked transcription units, showing genes sets with q values of <0.05, plotted against % genes in overlap. Gene sets significantly affected include those linked with breast cancer, and those regulated by polycomb and TGFβ.

### SMAC integrity and de-ubiquitylase access

Disruption of SMACs by DNF’s could deplete H2AK119ub1 in Xi chromatin by failing to support recruitment or concentration of the catalytic activity of PRC1 (RING1A/B), or conversely by allowing access to de-ubiquitinating enzymes. BAP1 is the catalytic subunit of the deubiquitinating enzyme complex (DUB) that removes PRC1-mediated H2AK119ub1, acting to restrict its deposition to polycomb target sites ^26^. To distinguish between recruitment and barrier functions (Fig.5C) we used PR-619, a broad-spectrum reversible inhibitor of DUB’s, including the PR-DUB BAP1^27^.

In transient transfection experiments the immediate (within 24 hours) loss of H2AK119ub1 from Xi was significantly blocked by PR-619 (SFig.7B), and in longer (3 day) transduction experiments the same trend was observed (Fig.5D), despite efficient SMAC dispersal by C181. Moreover, even in genetically CIZ1 null PEFs in which H2AK119ub1 is typically absent from Xi chromatin ^8^, its enrichment (but not that of H3K27me3) was restored within 24 hours of exposure to PR-619 (Fig.5E). Thus, loss of CIZ1 SMACs, whether by genetic deletion or dispersal by DNF’s, supresses accumulation of H2AK119ub1 in Xi chromatin in a manner dependent on DUB activity. We suggest therefore that CIZ1 SMACs perform a shield function that protects Xi chromatin from enzymatic attack by DUBs.

It should be noted here that in 3 day lentiviral transduction experiments with C181, we also measured a nucleus-wide *increase* in H2AK119ub1 levels in both of the WT female PEF populations analysed (SFig.8A). This was accompanied by changes in the relative levels of RING1A and RING1B protein (SFig.8B), both of which can support H2A ubiquitination on the inactive X chromosome ^28, 29^, but which may have different modes of target selection analogous to that described for EZH2 variants ^8^. These observations raise the possibility that loss of SMAC integrity, and associated exposure to DUBs, drives compensatory changes in PRC1.

### CIZ1 dominant negative fragments alter gene expression

To confirm that CIZ1 DNF’s have the potential to affect gene expression, and therefore eventually cellular identity, we analysed transcriptomes of three independent populations of early passage WT PEFs, isolated from two female and one male embryo, after transduction with C181 or with empty lentiviral vector for 3 days (Fig.5A), and also compared the effect to non-transduced cells (SFig.8C). Consistent changes in 510 transcription units (log2 fold change >0.58, FDR q-value 0.05) were observed (Fig.5F, SFig.8D, Supplemental Data set 2), of which 464 were named coding genes. Gene set enrichment analysis revealed significant similarity with 50 molecular signatures derived by chemical or genetic perturbation in murine cells (GSEA MSig. mCGP), of which the most significant are those linked with the developmental regulator transforming growth factor beta (Fig.5G, SFig.8E). Also significant is the polycomb complex-related gene set targets of EZH2 (M1486), and three gene sets related to breast cancer or mammary stem cell phenotype (M2573, M9908, M15150, SFig.8F). This shows that, similar to deletion of CIZ1, interference with CIZ1 assemblies can consistently and significantly alter gene expression, including genes linked with cellular plasticity and cancer. Moreover, as in experiments where CIZ1 is reintroduced against a CIZ1 null background ^8^, the effect is rapid (within days) and coincident with changes in chromatin. Together the data argue for a potent effect of CIZ1 DNFs, that results in destabilisation of the nucleus.

## Discussion

The primary purpose of heterochromatin formation during development is to protect and reinforce cell fate decisions, by restricting transcription factor access to genes. Thus, potential stabilizers of chromatin state, whose mis-expression may drive heterochromatin instability, are important to understand in relation to degeneration of cellular identity, human disease and aging. Our data suggest that RNA-dependent CIZ1 assemblies, exemplified by the *Xist*- and PLD-dependent CIZ1 SMACs that surround the inactive X chromosome, normally act as a molecular shield that helps protect Xi heterochromatin from the action of PR-DUBs.

‘Molecular shield’ is one of eight functional classes proposed for phase separating proteins outlined by PhaSePro ^30^. Defined as membrane-less organelles that inactivate reactions by sequestering some of the required components while keeping others outside, a CIZ1 shield would in this case sequester chromatin while excluding PR-DUBs. Questions remain about the structure and influence of such a shield. Does it impart a sealed barrier function, or a field through which some molecules pass more freely than others? We present some clues here about the structure of CIZ1 assemblies - apparently made up of aggregates of antiparallel homodimers that interface through an MH3 domain - but much remains to be uncovered about their structure, permeability and stability.

The polycomb repressive complex (PRC) family of proteins responsible for H2AK119ub1 and H3K27me3, and the PR-DUBs that remove ubiquitin, are already deeply implicated in human disease states, particularly cancer, and interfering with their function offers potentially powerful anti-cancer strategies ^31, 32^. The hitherto unappreciated barrier created by CIZ1 assemblies has the potential to locally buffer the activity of PR-DUBs, and so tip the balance between ubiquitination of H2AK119 and its removal. Here, we extrapolate to suggest that cells with intact shield function might be less sensitive to PR-DUB inhibitors than those in which it is dissipated, with favourable effects on cancer selectivity.

### Mechanism of DNF action

Exclusion of the N-terminal RD domain, which encodes the two PLDs that confer ability to coalesce inside the nucleus ^9^, converts CIZ1 from a SMAC participant into a molecule with the ability to disperse SMACs – a SMAC buster. Our functional modelling suggests that dominant-negative dispersal activity reflects retained ability to dimerise through its MH3 domain, but reduced ability of partial dimers to coalesce. Thus, the shift towards expression of truncated CIZ1 variants in human tumours would be predicted to interfere with normal CIZ1 function in heterochromatin protection, and so contribute to decay of the epigenetic landscape. By analogy with the way in which loss-of DNA repair functions propel cells toward genetic instability, our data suggest that CIZ1 DNF’s may propel cells toward epigenetic instability.

How might DNF-driven shield failure contribute to genetic heterogeneity and cancer aetiology? Evidence from several sources supports the proposition that disruption of polycomb function may precede genetic change. For example, PRC1-dependent H2AK119 ubiquitylation is implicated in genome integrity through recruitment of DNA repair factors during the DNA damage response ^33^, which acts as a barrier to genomic instability and cancer ^34^. Similarly, progression of uveal melanoma is thought to be driven by loss of PRC1, in this case through dysregulation of target genes linked with mitotic chromosome instability ^35^. Thus, established routes exist through which the observed expression of DNFs in early stage cancers could contribute to tumour initiation.

While the data presented here shows de-protection of Xi chromatin and erosion of H2AK119ub1 enrichment at Xi, the effect of CIZ1 DNF’s may extend beyond the Xi, and does extend beyond H2AK119ub1 to H3K27me3. In addition, at least one other histone post-translational modification, H4K20me1 (added by Set7/Set8/KMT5a), is depleted from Xi chromatin in cells that lack CIZ1 ^36^. This suggests that the effect of DNFs may extend beyond disruption of the polycomb axis.

### Dysregulation of CIZ1 expression

Our proposal is that expression of DNFs precedes disruption of chromatin state, which precedes widespread genetic instability. This poses the obvious question of what drives DNF expression? There are no recurrent polymorphisms in CIZ1 in adult cancers, and only one part-copy number variation is reported in COSMIC across 40562 unique samples. Therefore, CIZ1 is not recognised as a ‘cancer’ gene ^37^. This contrasts with chromatin proteins and their modifiers in which mutations occur in approximately half of all tumours ^38^. Despite this, and despite no overall changes in CIZ1 levels (TCGA transcripts averaged across the whole gene), CIZ1 dysregulation has been reported for a range of common solid tumours including breast ^39^, colorectal ^40, 41^, gallbladder ^42^, liver ^43^, lung ^44^ and prostate ^45^, mostly by sampling with site-specific detection tools. Thus cancer-associated, and potentially cancer-driving, changes in CIZ1 have not yet been captured by large-scale genome or transcriptome studies.

Lack of mutations raises the possibility that, not only can CIZ1 exert an effect epigenetically, it may itself be perturbed primarily epigenetically with DNF expression driven by as yet unknown triggers. The forms observed here, evidenced by transcript elevation from exon 10, are also detected in non-diseased cells but to a lesser extent. This suggests a normal biological context, perhaps to confer fluidity on SMACs as cells pass through natural transition states. A pause or delay imposed by extrinsic conditions might therefore prolong residency, and so exposure to the destabilizing effect of DNFs. Alternatively, over production of DNFs could be driven more directly, by dysregulation of transcription or transcript stability at some level. At present the question of what drives DNF expression in early stage breast cancers remains unanswered.

### Concluding lines

There remain fundamental questions about the relationship between genetic and epigenetic models of cancer and the question of which comes first is likely to have a range of context specific answers. For some types of tumour no genetic driver mutations are detected ^46^, and transient loss of PRC proteins appears to be sufficient to initiate cancer phenotypes in experimental contexts ^47^. Expression of CIZ1 DNFs in early stage breast cancers, and their ability to shift the polycomb axis, is therefore a possible causative event. However, based on the available functional information, products of the CIZ1 gene can be both epigenetic stabiliser and so candidate tumour suppressor (SMAC builder), and also destabiliser and so candidate tumour driver (SMAC buster). Assuming it were possible to selectively supress SMAC buster but not SMAC builder, a key question is whether intervention could have a clinical impact in developed tumours that have gone on to acquire genetic change. Perhaps a more realistic, and exciting, strategy would be to supress CIZ1 DNFs with the aim of dampening pre-genetic instability in order to promote cancer avoidance.

## Author contributions

GT, EL, ES, LB, SS, JA, AAA, AM, DC designed experiments. GT, EL, ES, LB, HS, DC performed experiments. SS, AA contributed to preparation of materials. GT and DC wrote the paper.

## Acknowledgments.

We thank Andrew Leach and Adam Dowle for specialist help with protein analysis, Will Brackenbury and Jonathan Godwin for cells, Christian Fermer of FDAB for anti-CIZ1 antibodies, and present and former colleagues Faisal Abdel Rahman, Maria Chechik, Jennifer Munkley, Louisa Williamson, Gillian Higgins, Julie Tucker, and Matt Dowson. We acknowledge the role of the Breast Cancer Now Tissue Bank in collecting and making available the primary normal breast epithelial cells used here, and the patients who donated their tissues. Protein domain analysis was funded by the Georgina Gatenby PhD scholarship to GT, human CIZ1 gene expression work by Cizzle Biotech, and other work by MR/V029088 and a Royal Society Leverhulme Trust Fellowship to DC.

## Methods

### Materials availability and contacts

Plasmids generated in this study [will be] deposited to Addgene, or will be made available on request. Further information and requests for resources and reagents should be directed to the lead contacts, Gabrielle Turvey and Dawn Coverley.

### Human Primary Cells

Human mammary epithelial cells (HMECs) were cultured at 37°C with 5% CO_2_ in MEBM basal medium (Lonza) supplemented with MEGM SingleQuots (Lonza) as per manufacturer instructions, on culture dishes coated in collagen (Thistle Scientific) as per manufacturer’s instructions to support their growth. Cells were sampled after limited passaging (p1-p2). Primary human mammary epithelial cells were acquired with informed consent from three donors by Breast Cancer Now Tissue bank under NHS ethical approval, and accessed under local approval from the University of York Department of Biology Research Ethics Committee.

### Human Cell Lines

All cell lines used are of female origin and were authenticated for this study by Eurofins Genomics human cell line authentication service (Eurofins Medigenomix Forensik GmbH) and results interpreted using CLASTR. Raw data is shown in STable 2 which returns the expected identities with 92-100% confidence in all cases. MCF-10A is a non-tumorigenic epithelial cell line established from human mammary gland with fibrocystic disease. MCF7 is a poorly-aggressive and non-invasive triple receptor positive human breast cancer cell line established from epithelial cells isolated from a metastatic mammary adenocarcinoma. BT-474 is a human breast cancer cell line established from a malignant ductal carcinoma of the breast that overexpresses human epidermal growth factors receptors 2 (HER-2) and oestrogen receptors (ER). SK-BR-3 is a human breast cancer cell line established from a malignant adenocarcinoma of the breast that overexpresses HER-2. MDA-MB-231 is a human breast epithelial cancer cell line established from a metastatic poorly differentiated triple-negative mammary adenocarcinoma. Cells were cultured in the following media: MCF-10A – MEGM, 5% horse serum,10 µg/mL hydrocortisone, 20 ng/mL EGF, 500 ng/mL insulin, 100 ng/mL cholera toxin, 1% PSG; MCF7 – EMEM, 10% foetal bovine serum (FBS) (PAA gold), 1% Penicillin-Streptomycin-Glutamine (PSG) (Gibco); BT-474 and SK-BR-3 – DMEM, 10% FBS, 1% PSG; MDA-MB-231 – DMEM, 5% FBS, 1% PSG.

### Mouse Primary Cells

All mouse primary embryonic fibroblast (PEF) strains (WT 13.24, 13.31, 13.32, 13.33, 13.27, 45.1fc and CIZ1 null 13.17, 41.2fa) were derived from day 13 embryos from C57BL/6 mice as previously described ^6, 8^. CIZ1 null mice were generated from C57BL/6 ES clone IST13830B6 (TIGM) harbouring a neomycin resistance gene trap inserted downstream of exon 1. The absence of *CIZ1*/CIZ1 in homozygous progeny was confirmed by qPCR, immunofluorescence and immunoblot.

All work with animal models was compliant with UK ethical regulations. Breeding of mice was carried out under UK Home Office license and with approval of the Animal Welfare and Ethical Review Body at the University of York. PEFs were cultured in 4.5g/L glucose DMEM containing 10% FBS, 1% PSG up to a maximum of passage 3. After passage 4 these cells are referred to as MEFs and were not used here.

### Mouse Cell Line

The female D3T3 cell line was cultured as described ^8^ in DMEM (Sigma), 10% FBS, 1% PSG (Gibco).

### Bacterial expression and purification of protein fragments

Murine CIZ1 constructs with an in-frame N-terminal glutathione S-transferase (GST) tag in pGEX-6P expression plasmids were expressed in BL21-CodonPlus-RP *E. coli* using lactose-driven auto-induction. A starter culture of bacteria was incubated in LB broth (Merck) containing selective antibiotic overnight at 37°C. Autoinduction media (1% w/v tryptone, 0.5% w/v yeast extract, 25 mM (NH_4_)2SO_4_, 50 mM KH_2_PO_4_, 50 mM Na_2_HPO_4_, 0.5% glycerol, 0.2% lactose, 0.05% glucose, 1 mM MgSO_4_, 1mM CaCl_2_ and trace metals: 50µM FeCl_3_, 10µM MnCl_2_, 10µM ZnSO_4_, 2µM CoCl_2_, 2µM CuCl_2_, 2µM NiCl_2_, 2µM Na_2_MoO_4_, 2µM Na_2_SeO_3_, 2µM H_3_BO_3_) containing selective antibiotic, was inoculated with starter culture until the OD_600_ was between 0.1-0.15 and incubated at 20°C for 24 h. Centrifugation in a Sorvall LYNX 6000 with a F9-6×1000 LEX rotor (4392 xg, 4°C, 15 min), produced a bacterial pellet which was resuspended in PBS buffer (137 mM NaCl, 8.1 mM Na_2_HPO_4_, 2.7 mM KCl, 1.5 mM KH_2_PO_4_) supplemented with 1 in 100 EZBlock™ protease inhibitor cocktail (BioVision) and 1 mM PMSF. Bacterial cells were lysed on ice via sonication for 5 cycles (30s on, 30s off) at 60% amplitude using 6mm probe (microtip MS 73, Bandelin SONOPULS). Lysates were clarified by centrifugation (25000 g 4°C, 30 min) in a Heraeus™ Multifuge™ X1 centrifuge with a F15-6×100y fixed angle rotor. All chromatography was performed on an ÄKTA™ chromatography system. Lysates were loaded onto a 5 mL glutathione sepharose column (Cytiva) at 0.5 mL/min, then washed extensively with 10 column volumes of PBS buffer followed by 10 column volumes of cleavage buffer (50 mM Tris-HCl, 7.0, 150 mM NaCl, 1 mM EDTA, 1 mM Dithiothreitol, pH 7), both at 1 mL/min. This was incubated at 4°C overnight with 2 units of PreScission protease (GE Healthcare) in cleavage buffer. Cleaved protein was eluted in fresh cleavage buffer and concentrated to 0.5 mL using a centrifugal concentrator (Sartorius). Protein concentration and quality was determined by NanoDrop® ND-1000 spectrophotometer (Labtech, version V3.2.1). Purified protein was supplemented to 5% v/v glycerol and snap frozen in liquid nitrogen and stored at -80°C.

### SEC and SEC-MALLs analysis

Proteins were concentrated to a final volume of 0.5 mL and 0.45µm filtered (Corning ®) prior to size exclusion chromatography (SEC). Total sample between 2-5 mg/mL was injected. Experiments were conducted at room temperature on an ÄKTApurifier™ system with Superdex 200 increase 10/300 GL column, equilibrated with 2 column volumes of running buffer before use. Chromatography was conducted at 0.5 mL/min flow rate using running buffer (50 mM Tris-HCl, 7.0, 150 mM NaCl, 1 mM EDTA, 1 mM Dithiothreitol, pH 7), and output monitored with a UV_280_ detector. Blank buffer injections were used where appropriate to prevent carry-over between sample runs. Samples were collected on the Frac-950 fraction collection system. Data was analysed using the UNICORN™ control software.

Size exclusion chromatography multiple laser light scattering (SEC-MALLs) analysis was used to determine the molecular weight of CIZ1 forms. 100 μl of sample between 2-5 mg/mL was injected at 0.5 mL/min flow rate at room temperature. The system was comprised of a Wyatt HELEOS-II multi-angle light scattering detector and a Wyatt rEX refractive index detector linked to a Shimadzu HPLC system (SPD-20A UV detector, LC20-AD isocratic pump system, DGU-20A3 degasser and SIL-20A autosampler). Solvent was 0.2 µm filtered before use and a further 0.1 µm filter was present in the flow path. The Superdex S200 10/300 GL column was equilibrated with at least 2 column volumes of running buffer before use, and flow was continued at the working flow rate until baselines for UV, light scattering and refractive index detectors were all stable. Shimadzu LCsolution software was used to control the HPLC and Astra V software for the HELEOS-II and rEX detectors. Blank buffer injections were used as appropriate to check for carry-over between sample runs. Data was analysed using Astra V software. Molecular weights were estimated using the Zimm fit method with degree 1. A value of 0.19 was used for protein refractive index increment (dn/dc).

### Electrophoretic mobility shift assay

EMSAs using C181 and deletion mutant lacking the MH3 domain, were carried out as previously described ^9^.

### RNA and Protein Extraction from mammalian cells

Total RNA or whole cell protein lysates were isolated from 9 cm culture plates at approximately 80% confluence. RNA was isolated using 2 mL TRIzol reagent (Invitrogen) as recommended, and protein lysates were scrape harvested directly into SDS-PAGE sample loading buffer (final concentration 75 mM Tris pH 6.8, 4% w/v SDS, 15% v/v glycerol, 250 mM 2-Mercaptoethanol) supplemented with 1 mM PMSF.

### Western Blotting

Samples (purified proteins or cell lysates) were denatured in gel loading buffer (final concentration: 4% SDS w/v, 75 mM Tris pH 6.8, 15% v/v glycerol, 250 mM 2-mercaptoethanol, bromophenol blue dye) at 95°C for 5 min, vortexed, heated for an additional 5 min, and centrifuged prior to loading. Samples were electrophoresed through 4-15% Mini-PROTEAN® TGX™ precast acrylamide gels (Bio-Rad) submerged in running buffer (250 mM Tris, 1.92M Glycine, 1% w/v SDS) at 40V for 15 min, then 90V for approximately 1.5 h. Resolved proteins were transferred to nitrocellulose membrane (0.1 μM NC, Cytiva Amersham™ Protran™) using a semi-dry transfer cell in transfer buffer (300 mM Tris, 10 mM CAPS, 0.02% w/v SDS, 10% v/v methanol). For peptide mapping, dot blots were generated by directly spotting peptides (STable 4) in PBS onto nitrocellulose membrane. Membranes were blocked in 10% w/v milk in PBS containing 0.1% TWEEN®20 (PBST). Primary antibody was applied overnight at 4°C or for 1-2 h at RT, antibodies are listed in STable 3. Proteins were visualised using Pierce™ ECL Western Blotting Substrate (Thermo Scientific™) and processed using the PXi gel imaging system (Syngene).

### Site-directed mutagenesis

Mutagenic primers that contain additions, substitutions or deletions of murine CIZ1 were created for use in PCR mutagenesis, and are listed in STable 5. All primers were purchased at 0.025µmol (Merck) and 1mM stock solutions prepared in TE (10 mM Tris, 1 mM EDTA) and stored at -20°C. Each PCR reaction was composed of CloneAmp HiFi PCR premix (Takara), 4ng of the plasmid template and 10-20ng of the primer pair mix, to a 10 µl total volume. PCR cycling conditions were: initial denaturation 98°C for 30 s; 18 cycles at 98°C for 10 s, 55°C for 30 s, 72°C for 15 s/kb; final extension 72°C for 10 s. Confirmation of PCR reaction was carried out on a 1% w/v agarose in 1x TBE (89 mM tris, 89 mM boric acid, 2 mM EDTA) gel containing 1x SYBR™ Safe DNA gel stain (Invitrogen™) using 5 μl of the reaction with 1 μl of loading dye (New England Biolabs). Agarose gel was electrophoresed at 70V until resolution of bands required was reached. Gels were visualised using a transilluminator in the PXi gel imaging system (Syngene). The unedited methylated DNA template present in the remaining sample was digested for 1 h at 37°C with DpnI restriction enzyme (New England Biolabs) in CutSmart® Buffer (New England Biolabs), as per manufacturer’s recommendation. This was subsequently transformed into competent cells to repair DNA nicks in the mutant plasmid. During transformation 1 μl of plasmid PCR product was incubated with 25 μl of DH5α competent cells for 30 min on ice, cells underwent heat shock at 42°C for 45 s, and finally incubated on ice for 2 min. This was allowed to recover for 1-2 h, depending on the selective antibiotic used, in 225 μl SOC broth (Invitrogen™) at 37°C in an orbital shaker at 200rpm. Bacteria were then plated onto antibiotic selective LB agar at varying concentrations for optimum colony production and incubated overnight at 37°C. Individual colonies were selected and grown in 5 ml antibiotic selective media overnight. The plasmid DNA was extracted using QIAprep Spin Miniprep Kit (QIAGEN), as per manufacturer’s instructions. All plasmids were sequence verified to confirm mutations (Eurofins TubeSeq Service). If the mutant was designed for protein production in bacteria this was additionally transformed into BL21-CodonPlus-RP *E. coli*.

### Lentivirus transduction

Bicistronic ZsGreen/C181-bearing virus and ZsGreen alone bearing virus were produced in the Lenti-X™ 293T subclone of human embryonic kidney (HEK) cells. 8×10^5^ HEK cells were seeded per well in a 6 well plate prior to transfection with plasmids. For transfection of each well, 1 µg transfer vector, 0.75 µg packaging plasmid, 0.25 µg envelope plasmid diluted in 100 µl optiMEM (Gibco) was mixed with 20 µl of PolyFect transfection reagent (Qiagen) and incubated for 5-10 min at room temperature to allow complex formation. 0.6 ml of cell growth medium was added and gently mixed then transferred to one well. Cells were incubated for 16 h, then media was replaced with fresh growth medium (supplemented with addition of HEPES to a final concentration of 20 mM). At 48 h post-transfection the supernatant containing virus was harvested and filtered through a low-protein binding filter (0.45µm, Sarstedt) to remove HEK debris. Viral supernatant was supplemented with 4 μg/ml polybrene (Sigma), and transferred to recipient PEFs or D3T3 cells. Transduction was monitored by emergence of cytoplasmic ZsGreen, and showed that close to 100% of the cells were transduced after approximately 48h.

### Transient Transfection

For analysis in cycling cells, cells were seeded on 13mm glass coverslips (SLS) at approximately 30% confluency one day prior to transfection, to produce populations at ∼60% confluency at time of transfection. Coverslips were transferred to individual wells in 24 well plate in 500μl media prior to transfection. For each coverslip 50 μl Opti-MEM® Medium (Gibco) was mixed with 1.5 μl X2 Transfection Reagent (Mirus) and 200ng plasmid DNA (pEGFP-C2 containing an insert derived from CIZ1), incubated for 30 min, then applied to cells dropwise. Coverslips were fixed and processed for immunofluorescence 24-48 hrs later. For contact inhibited cells, cells were plated across a range of densities by serial dilution two days prior to transfection. Coverslips at greater than 90% confluency were selected for transfection, and processed as above.

### Cell synchrony

D3T3 cells were arrested in mitosis or S phase by application of 50 ng/ml nocodazole (Sigma) for 16-24 h, or 2.5 mM thymidine (Sigma) for 24 h, respectively. Cells arrested in M phase were isolated by mitotic shake off and re-plated onto glass coverslips for analysis post release. Cells held in S phase grown on glass coverslips were released by washing twice with PBS, then replacing with fresh media. In transduced cell populations, cells were arrested approximately 48 hrs post transduction for 16-24 hr, then released and analysed. To facilitate retention of mitotic cells, coverslips were fixed prior to permeabilization.

### Flow cytometry

Cells were isolated from 9 cm culture plates by trypsinization and resuspended in 100 μl cold PBS to obtain a single cell suspension, then stored at -20°C after addition of 1.5 ml cold 70% ethanol. For analysis cells were pelleted and resuspended in PBS (500,000 cells/mL), and 55 μl 10x FACS mix (1 mg/ml propidium iodide, 4% v/v Triton X-100, 10xPBS) was added per 500 μl of cell suspension. DNA content was measured using CytoFLEX (Beckman Coulter) at excitation 561 nm/emission 610/20 for detection of the DNA binding dye propidium iodide ^48^. A minimum of 5,000 single cells per sample were recorded for analysis using cell cycle algorithm software FCS Express V7 (Dotmatics).

### Inhibitors

To measure the impact of inhibition of PR-DUBs, 5µM PR-619 (Bio-Techne) was applied to PEFs 16 h post transduction for 32 hrs, to collect cells 48 h post transduction. In transient transfection experiments PR-619 was used at 5 µM for 24 h during the transfection window.

### Immunofluorescence

Cells grown on coverslips were washed in cytoskeletal buffer (10 mM PIPES/KOH pH 6.8, 100 mM NaCl, 300 mM sucrose, 1 mM EGTA, 1 mM MgCl_2_) with 0.1% v/v Triton X-100, and fixed in 4% w/v paraformaldehyde (PFA). Where indicated Triton X-100 was left out of CSK (unextracted cells), or additional 400mM NaCl was added (high-salt extraction). After fixation all coverslips were blocked in antibody buffer (AB) (1xPBS, 10 mg/ml BSA, 0.02% w/v SDS, and 0.1% v/v Triton X-100) for 30 min, incubated with primary antibodies for 1 h at 37°C, washed three times with AB, incubated with secondary antibodies for 1 h at 37°C, washed three times with AB, and mounted on glass slides with Vectashield mounting medium containing DAPI (Vector Labs). Primary antibodies are detailed in STable 3. Anti-human CIZ1 monoclonal antibodies 79 and 87 were generated by Fujirebio Diagnostic Antibodies (FDAB). Anti-species antibodies (ThermoFisher) labelled with Alexa Fluor 568 (red) or 488 (green) was used for detection in all cases. Fluorescence images were captured using a Zeiss Axiovert 200M fitted with a 63×/1.40 Plan-Apochromat objective and Zeiss filter sets 2, 10, and 15 (G365 FT395 LP420, BP450-490 FT510 BP515-565, and BP546/12 FT580 LP590), using Axiocam 506 mono and Axiovision image acquisition software (SE64 release 4.9.1) through Zeiss Immersol 518F. For each antibody constant image capture parameters were used to generate image sets within an experiment, on which quantitative analysis was performed, in all cases from unmodified raw images.

### Phenotype Scoring in Dispersal Assay

Cells were inspected by eye across replicate experiments, and across 2-3 replicate coverslips per condition within an experiment. Avoidance of bias was achieved by verification by independent workers in all cases, and blinded analysis in some cases. In the two-tier scoring system cells were categorised as either having a CIZ1 Xi assembly or not, and in the three-tier scoring system by inclusion of an intermediate category in which CIZ1 assemblies were reduced or diffuse, or made up of locally dispersed particles. Empty vector (EV) was used as a negative control, and WT-C181 as positive control in experiments to test the effect of mutants.

### Image analysis in Dispersal Assay

For measurement of the effect of DN fragments on endogenous CIZ1 or histone PTMs at Xi, or nucleus wide, sets of images including test and controls samples, were processed in parallel and imaged with identical parameters in one sitting. All intensity measurements were conducted on unedited, unenhanced raw image sets. FIJI identified regions of interest (ROI) within Dapi stained fields of nuclei using auto-thresholding with Otsu setting to create a binary mask that defined nuclear perimeters. ROI’s were applied to antibody-detected fluorescence image layers, to generate intensity means, minimum, and maxima per ROI, and area of each nucleus/ROI. In female nuclei, to obtain a surrogate estimate of Xi-assembly intensity, means for a subnuclear circle area (excluding the Xi) was subtracted from the intensity means for whole individual nuclei. Where two or more data sets (for example C181/vector control pairs) from experiments performed on different days or with different PEF populations were combined, summary data was generated after normalization of values to the average of the control set in each case, which was set at 1. For reproduction, images were digitally enhanced to remove background fluorescence or increase brightness using FIJI. Identical manipulations were applied within an experiment, so that for example, the intensity of staining before and after extraction with high-salt buffer, or with and without transfection, is accurately represented.

### Transcriptomics

Primary murine embryonic fibroblasts 13.31, 13.32 (WT female) and 13.33 (WT male) were transduced with virus bearing either the empty pLVX-EF1a-IRES-ZsGreen1 vector (Takara), or the same plasmid expressing the coding sequence of C181 as described in lentivirus transduction. After 72 hrs, RNA was extracted with TRIzol (Ambion 15596-026) following manufacturer’s instructions from both transduced plates and untransduced controls. RNA pellets were resuspended in nuclease-free water. Isolated RNA was treated with DNase (Roche 04716728001) before quality analysis by agarose gel, NanoDrop spectrophotometer and Agilent 2100 Bioanalyzer. Library preparation and sequencing was undertaken by Biomarker Technologies (BMKGene), using the NEBNext UltraTM RNA Library Prep Kit for Illumina® (NEB, USA), with enrichment for mRNA using oligo(dT)-magnetic beads, followed by random fragmentation of enriched mRNA in fragmentation buffer. cDNA was synthesized using random primers followed by purification with AMPure XP beads, end repair, dA-tailing, adaptor ligation, PCR enrichment and further AMPure XP purification to select fragments within size range of 300-400 bp. Library quality was assessed using the Agilent 2100 Bioanalyzer system and sequenced using an Illumina platform, using paired-end sequencing to generate at least 15.31Gb clean data per sample, with minimum 93.16% of clean data, quality score of Q30. Low quality sequence reads and adaptor sequences were removed and the resulting high-quality sequence reads were aligned to version GRCm38_release 79 of the mouse genome using HISAT2 ^49^. Transcriptomes were assembled and gene expression quantified using StringTie ^50^ Differential gene expression analysis was performed by DESeq2 ^51^. Bioinformatic analysis and plots were generated from the average of transcripts per million plus one (TPM+1) values for each treatment condition to exclude very lowly expressed transcripts using Spyder (v.5.3.3) accessed by Anaconda Distribution (v.2.3.2). Scatter plots were generated using the pandas, numpy and matplotlib modules. Volcano plots were generated using the pandas and bioinfokit modules. Principle component analysis (PCA) plots were generated using the pandas, sklearn, seaborn and matplotlib modules. Gene set enrichment analysis (GSEA) was performed using the GSEAPY module, with genes preranked based on generation of a π value as calculated by multiplication of the log_2_FC of the average TPM by the -log_10_(q value).

### Patient and Cell Line Bioinformatics

Aligned RNA sequencing data for 1095 primary breast cancer samples from The Cancer Genome Atlas were accessed under dbGaP project 25297. Adjacent normal RNA sequencing data was available for 113 donors. Secondary metastatic breast samples were excluded from analysis. Data were downloaded using the Genomic Data Commons command line client v1.5.0. FASTQ files were regenerated from sample BAM files using samtools v1.10 ^52^ to exclude secondary and supplementary alignments, and then BEDTools v2.27.1 bamToFastq ^53^. No additional quality control steps were performed on the extracted read files. Reads were aligned to the GRCh38 Gencode primary assembly and to individual *CIZ1* transcripts (from Gencode v38 and ^54^) using HISAT2 v2.2.0 ^55^. Reads were also pseudo-aligned to the Gencode v38 full annotation transcriptome file with kallisto v0.46.0 ^56^, quantified and aggregated to gene-level transcripts per million (TPM) expression values using tximport v1.24.0 ^57^. The same expression analysis pipeline was also completed on publicly available RNA sequencing data from four breast cancer cell lines (MCF7: SRR8615758; BT-474: SRR8616195; SK-BR-3: SRR8615677; MDA-MB-231: SRR8615767) and one breast epithelium transformed cell lines (MCF-10A: SRR12877369). *CIZ1* transcript read coverage was normalised to the canonical (ENST00000372938.10) exon 7 coverage, and then stratified by tumour stage. Similar analysis of *ESR1* and *TP53* was completed to rule out broader 3’ coverage biases. *CIZ1* alignments from both transcript and full genome mappings were inspected manually for novel, well-supported splice junctions in IGV Desktop for Windows v2.8.2 ^58^. The top 10 stage II patients used to check for 3’ bias were: TCGA-E9-A54Y, TCGA-LL-A6FR, TCGA-E9-A3X8, TCGA-AQ-A54O, TCGA-AQ-A54N, TCGA-WT-AB41, TCGA-A2-A3XV, TCGA-LL-A5YL, TCGA-GM-A2DB, TCGA-AO-A03N.

### Gene Expression

To quantify relative expression of *CIZ1* amplicons in primary tumours, TissueScan_TM_ Tumour cDNA arrays from OriGene Technologies, Inc. (Rockville, MD) containing 2-3 ng of cDNA were analysed by qPCR. The isolated RNA used to generate the cDNA arrays was collected under IRB approved protocols, and array details with tumour classification is given in Supplemental Data set 1. Tumour classifications and abstracted pathology reports are given at: http://www.origene.com/qPCR/Tissue-qPCR-Arrays.aspx. cDNA was normalized using β-actin by the supplier, and we used our own amplification of β-actin where indicated. In most cases results for *CIZ1* amplicon expression were expressed relative to one another, rather than to another gene. Reactions were carried out in 25 µl volumes with 12.5 µl Taqman master mix (Applied Biosystems), 1 µl of each 10 µM primer and 1 µl 10 µM probe. Primers (Sigma Aldrich) and probes (MWG) are given in Stable Primers 9 and 10 were combined with a probe in exon 5 to generate detection tool set DT5, primers 13 and 14 with probe 7 (DT7), primers 1 and 2 with probe 14 (DT14), and primers 6 and 7 with probe 16 (DT16). Primer efficiencies were greater than 90% in all cases, and the relative amplification efficiencies of RD and AD tools were routinely checked using plasmid template with coupled and equal levels of RD and AD. Data was generated using an ABI 7000, SDS v1.2. (Applied Biosystems), using 50°C (2 min), 95°C (10 min), then 50 cycles of 95°C (15 s), 60°C (1 min). Relative expression was calculated using the comparative Ct method using the formula, 2^-ΔΔCt^ ^59^, and results expressed relative to the mean of normal cells or tissue in each array, or to the lowest stage tumour in the array, as indicated.

### Quantification And Statistical Analysis

For analysis of CIZ1 SMAC frequencies, a variable number of technical replicates and independent counts (N value) was conducted per experiment, allowing generation of ± SEM values as shown. The number of cells scored is stated individually in each experiment (n value). Wherever possible at least two independent PEF lines were used across the experimentation relating to each question. Statistical analysis was carried out in SPSS (IBM Corp. Released 2021. IBM SPSS Statistics for Macintosh, Version 28.0. Armonk, NY: IBM Corp) or Microsoft Excel, and parametric or non-parametric tests were utilised where appropriate. For comparison between two data sets a two-sample unpaired t-test or a Mann-Whitney U test was utilised. For comparison between three or more data sets a one-way ANOVA followed by an appropriate post-hoc test was utilised. Statistical tests used in each analysis are stated in the figure legend, alongside the p value generated. Graphs were generated using Microsoft Excel, and where shown, measurements normalised to the relevant control. Individual significance values are indicated, asterisks indicate *=p< 0.05, **=p< 0.01 and ***=p<0.001. For qRTPCR data Pearson correlation coefficient was used to compare CIZ1 RD and AD amplicon expression, and linear regression trendlines were applied using Excel ver.16.16.7. P values were generated by T-test in Excel.

## Materials

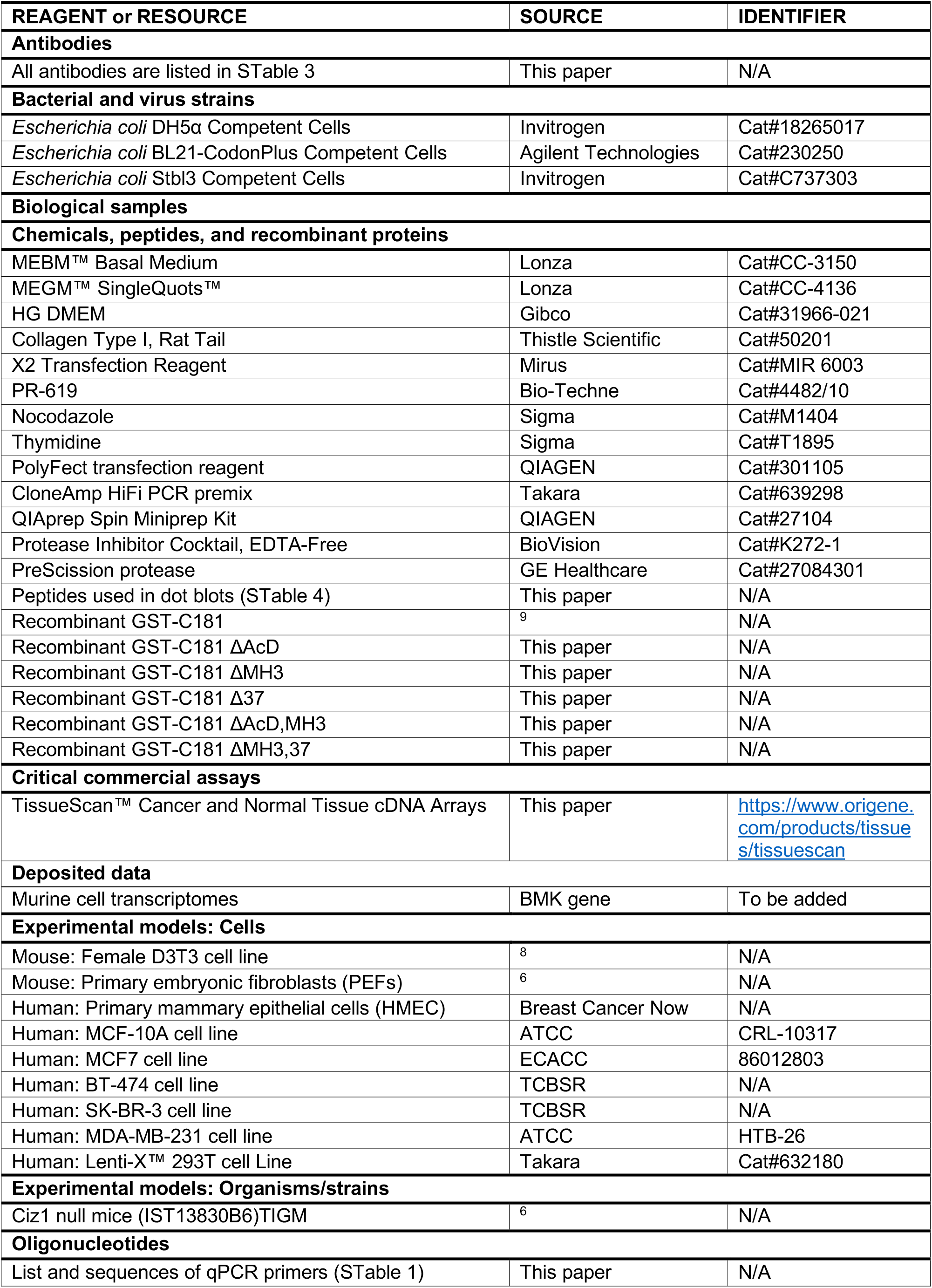
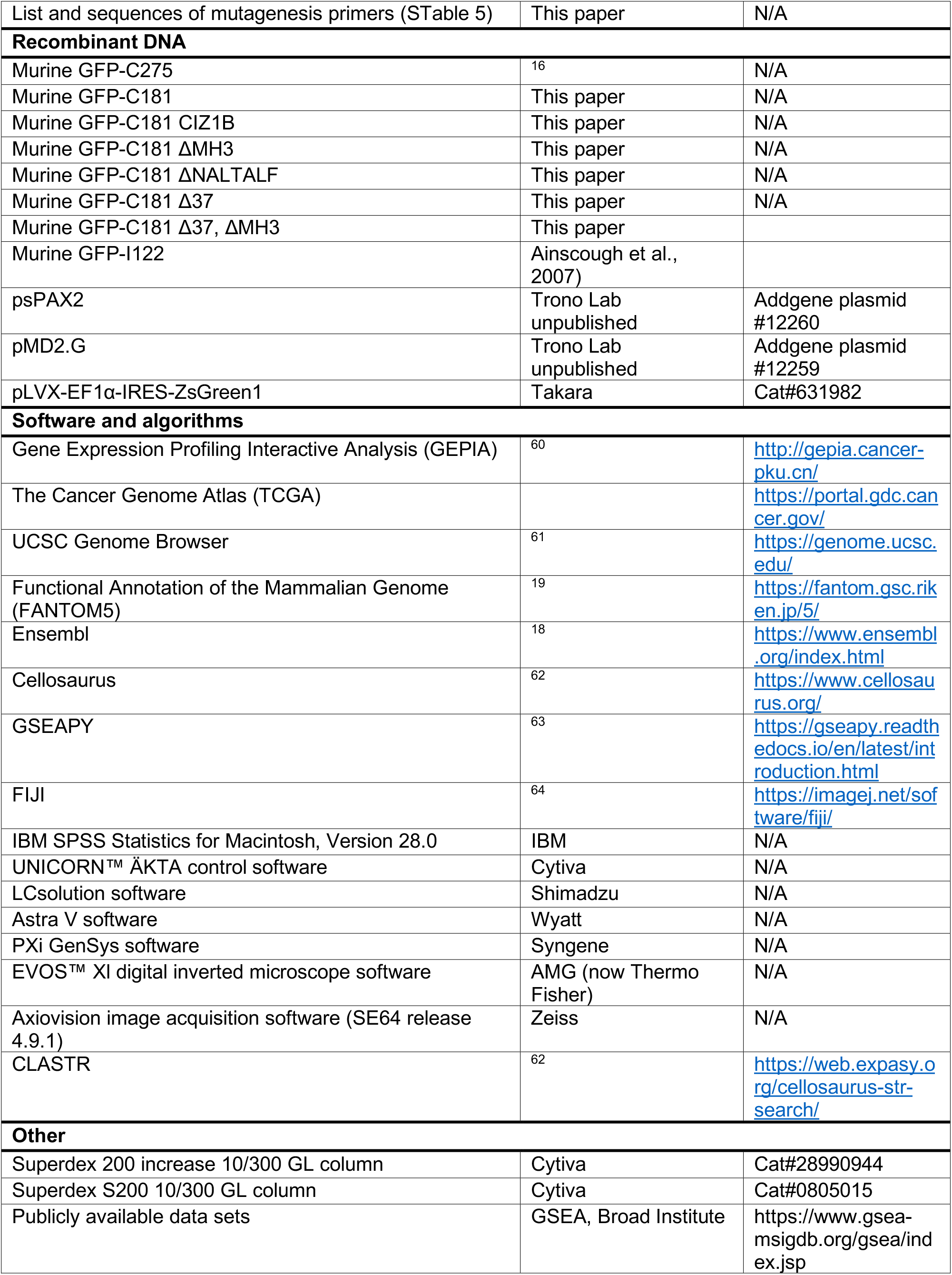

## Supplemental material

**Supplemental Figure 1.**
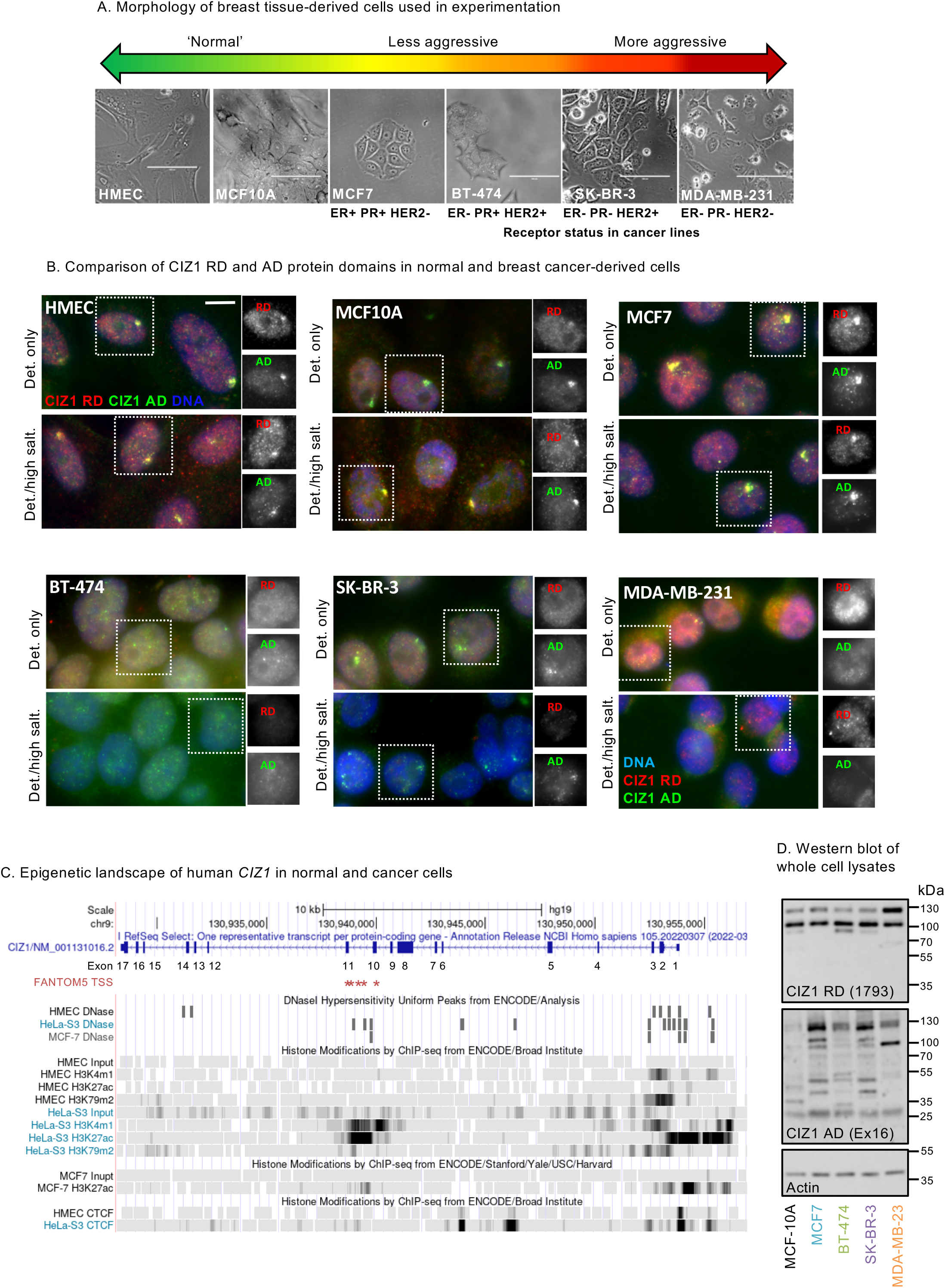
Corruption of CIZ1 Xi SMACs in breast cancer cell lines. A) Bright field images of breast-derived cell types ordered based on phenotype, with corresponding hormone and growth factor receptor status. Bar is 100 microns. B) As in Fig.1C, images of CIZ1-RD (red) and CIZ1-AD (green) after pre-fixation wash with detergent-containing buffer (Det. only), or after high-salt extraction (Det./high salt). Box shows nucleus for each cell type in which red and green stain are shown individually in grey scale. Bar is 10μm. The RD and AD epitopes were differentially extracted in some cases, indicating that they are not always part of the same polypeptide (for example compare nucleus-wide RD in SK-BR-3 cells, in detergent-treated cells to detergent/high-salt treated cells). C) *CIZ1* locus in *Homo sapiens* with corresponding exon numbers. Potential CIZ1 alternative transcription start sites (TSSs) in exons 10 and 11 predicted in the FANTOM5 project (Lizio et al., 2015) are indicated (red stars). The epigenetic landscape in human mammary epithelial cells (HMEC), a cervical cancer cell line (HeLa) and a breast cancer cell line (MCF7) is shown below. Diagram generated using UCSC genome browser (Kent et al., 2002). D) Western blots of whole cell lysates from all five cell lines analysed in Fig.1/SFig.1. N-terminal RD and C-terminal AD epitopes were detected by (Coverley et al., 2005), and rabbit pAb 329770 (Biorbyt), respectively.

**Supplemental Figure 2.**
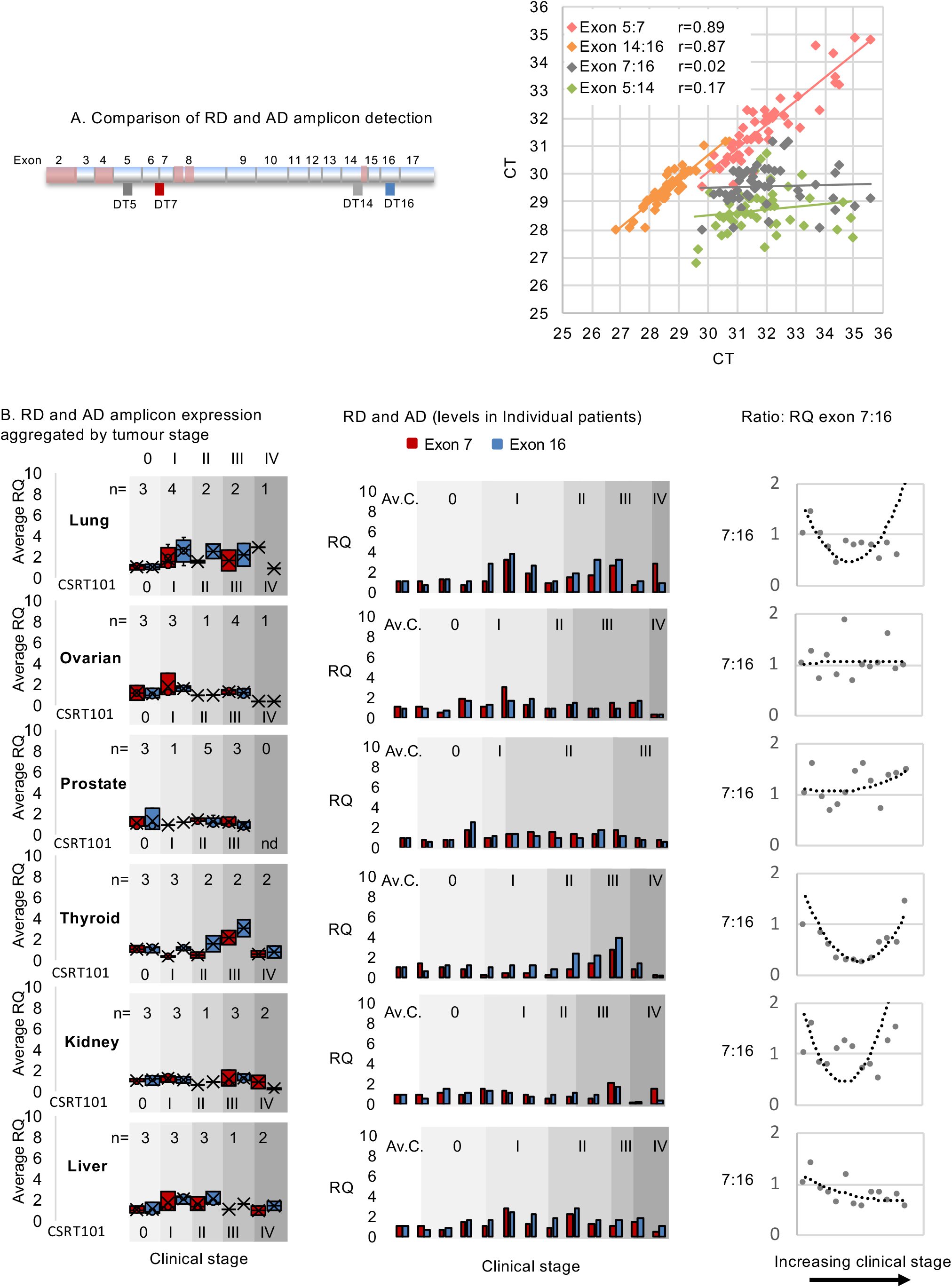
CIZ1 domain expression in common solid tumours. A) Exon structure of CIZ1 based on human reference sequence NM_012127.2 which encodes 898 amino acids, showing all 16 translated exons (2-17). Untranslated alternative exons 1’s are not shown. The boundaries of partially characterised alternatively-spliced exons are indicated in pink (Coverley et al., 2005; Dahmcke et al., 2008; Higgins et al., 2012; Rahman et al., 2007; Sofi et al., 2022; Swarts et al., 2018). The location of amplicons detected by human quantitative RT-PCR detection tools are also shown. These are four Taqman primer/probe sets; DT5 and DT7 which detect the 5’ end of CIZ1 transcripts, and DT14 and DT16 which detect the 3’ end (S.Table 1). Right, comparison of outputs with the indicated pairs of detection tools applied to 46 human tissue-derived cDNAs in Origene cDNA array. Pearson’s correlation coefficients show strong agreement between exons 5 and 7, and between 14 and 16, but poor agreement between exons 7 and 16, or 5 and 14. This shows that the 5’ and 3’ ends of CIZ1 are typically imbalanced at the transcript level. B) Left, relative expression of exons 7 (red) and 16 (blue), normalized to the average of 3 unmatched control samples for each of six common solid tumour types in multi-tissue cDNA array CSRT101 (Origene), aggregated by disease stage (0-IV, where 0 represents histologically normal tissue). Middle, individual tissue data plus the average of the controls calibrated to 1 (Av.C). Right, plots show the ratio of exon 7 to exon 16 ordered by stage, with trendline (polynomial 2). Individual sample information for all array samples in given in Supplemental Dataset 1.

**Supplemental Figure 3.**
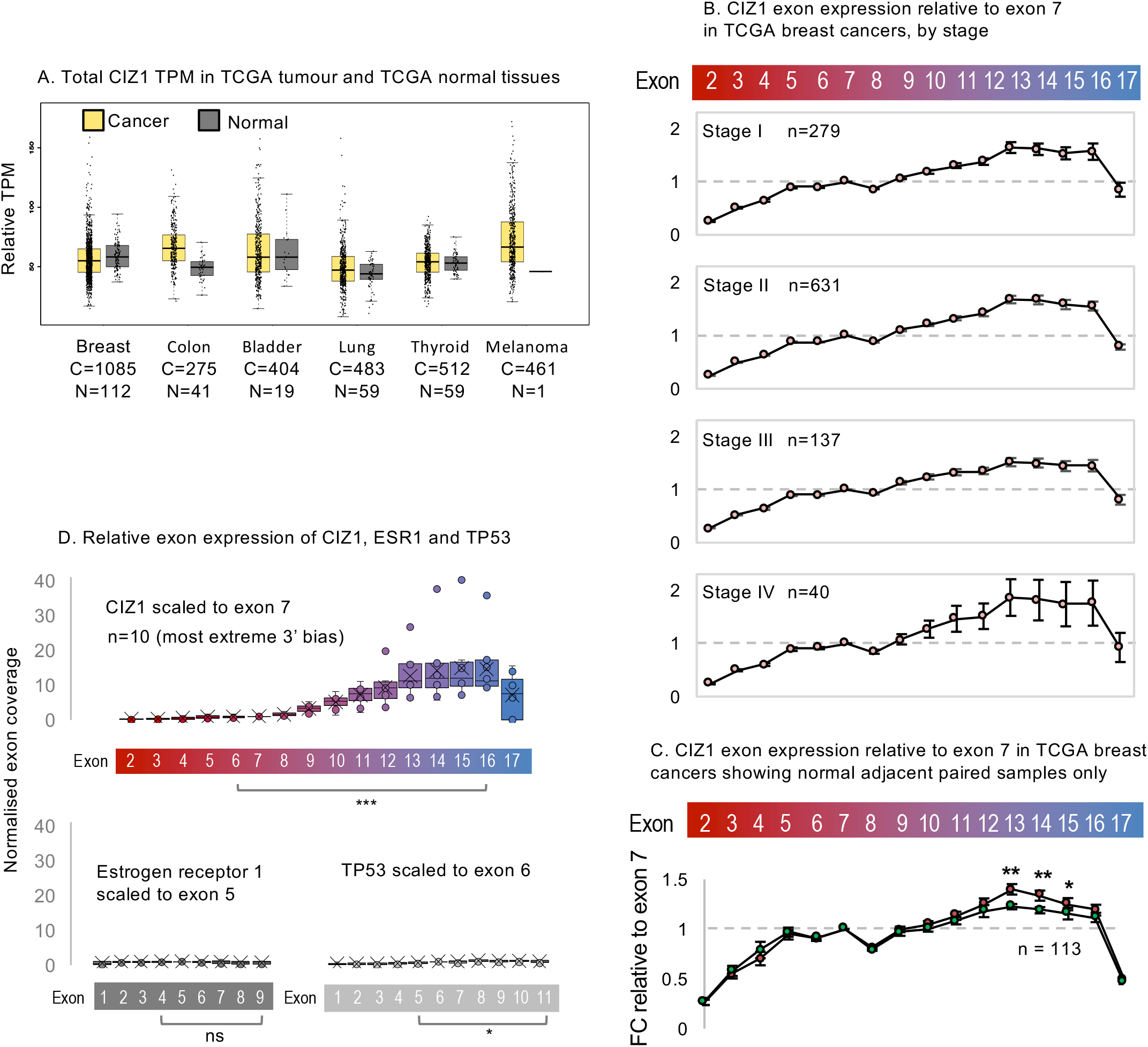
Extended analysis of CIZ1 in TCGA cancer patient transcriptomes. A) Total CIZ1 TPM derived from the indicated number of cancer (C) and normal (N) tissues in TCGA, compared using GEPIA http://gepia.cancer-pku.cn/, for the indicated disease types. No significant elevation is returned (where log_2_FC was >1, and p-value < 0.05), when comparing all amalgamated transcripts that map to the CIZ1 gene (not resolved by exon). B) All TCGA breast cancer-derived CIZ1 transcripts mapped to exon 2-17, normalized to each individual exon 7 as shown in Fig.2E, by stage. At all stages overrepresentation is first evident around exon 10, while exon 17 is under represented compared to 16. Error bars show SEM. C) TCGA breast tissue samples that have a matched paired normal adjacent tissue (n=113), showing CIZ1 transcripts mapped to exons 2-17, normalised to each individual exon 7. Comparison of exons is by Wilcoxon signed rank test. Breast cancer samples have a significant increase in expression of exon 13, 14 and 15 compared to their adjacent tissue (p=0.0013, p=0.0015 and p=0.025 respectively). D) Control analysis showing TPMs in a subset of 10 stage 2 TCGA breast cancer patients that exhibit the most marked 3’ end bias for CIZ1, mapped to CIZ1 exons, normalised to exon 7. Below, TPMs from the same patients for estrogen receptor alpha (ERα/ESR1) normalised to its exon 5, and TP53 normalised to its exon 6, showing relative exon coverage and lack of 3’ over-representation. Comparisons by t-test for the indicated exons, CIZ1 p=0.00045; ESR1 p=0.0504; TP54 p=0.037.

**Supplemental Figure 4.**
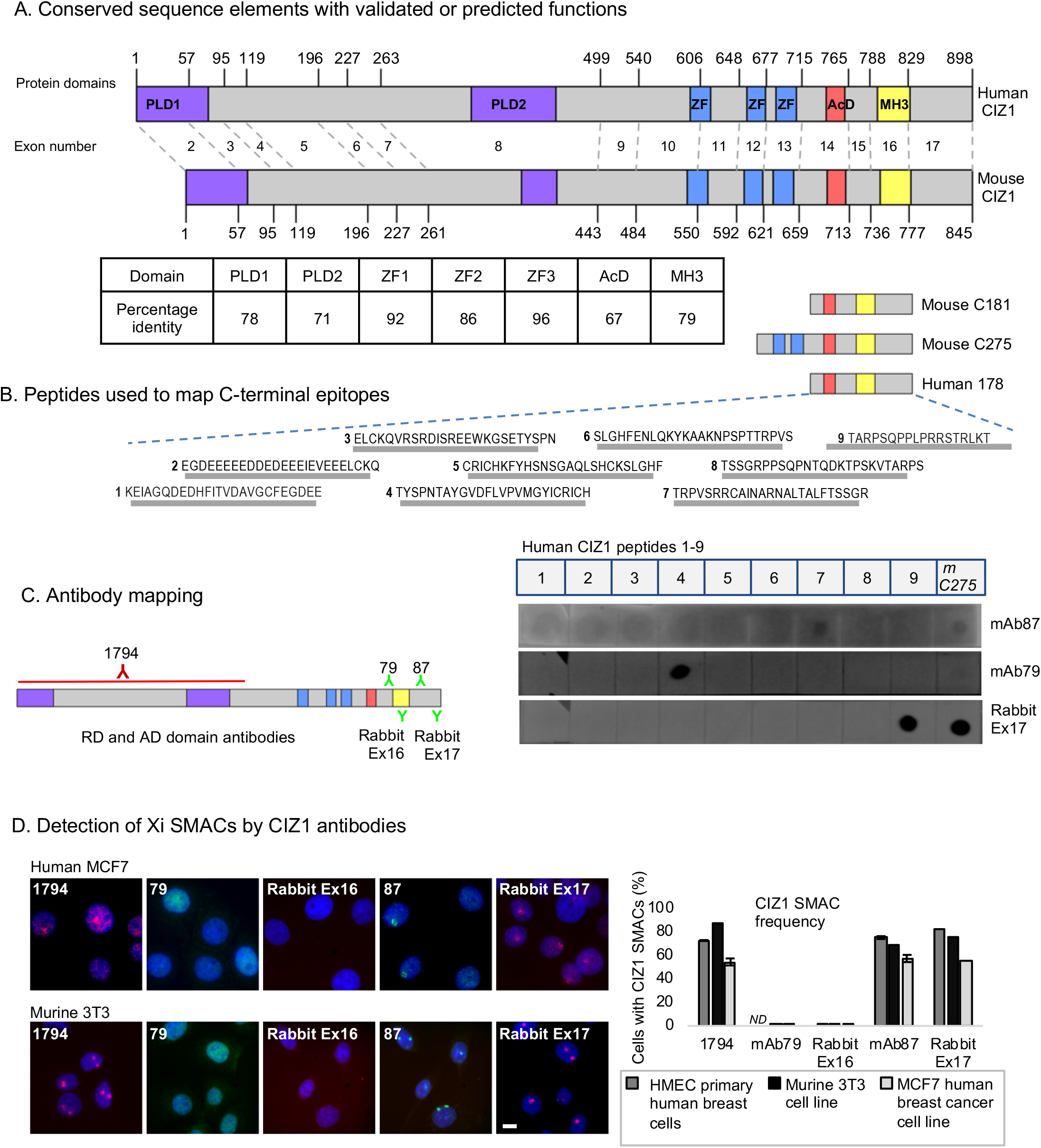
Antibodies, epitopes and protein domains in murine and human CIZ1. A) Protein domain map aligning human and mouse CIZ1. Numbers correspond to amino-acids encoded at exon boundaries. In human CIZ1 (reference sequence NP_001124488.1) the domains highlighted are: Prion like domains 1 and 2 (PLD1 and PLD2, purple) at position 1-78 and 360-451 respectively, three zinc fingers (ZF1, ZF2 and ZF3, blue) at positions 593-617, 656-676 and 687-709 respectively (ZnF_C2H2 SM00355, ZF_C2H2 sd00020 and ZF_C2H2 sd00020), an acidic domain (red) containing a highly concentrated area of aspartates and glutamates at position 741-761, and a matrin-3 homology domain (yellow) at position 796-831 (ZnF_U1 smart00451). In mouse CIZ1 (reference sequence NP_082688.1) the domains highlighted are: Prion like domain 1 and 2 (PLD1 and PLD2) at position 1-67 and 361-399 respectively (Sofi et al., 2022), three zinc fingers (ZF1, ZF2 and ZF3) at positions 537-561, 600-620 and 631-653 respectively (ZnF_C2H2 SM00355, ZF_C2H2 sd00020 and ZF_C2H2 sd00020), an acidic domain at position 689-709 and matrin-3 homology domain at position 746-770 (ZnF_U1 smart00451). Boxes show % identity at the amino-acid level across these domains. B) Synthetic peptides spanning human C-terminal 178 amino acids (C178), used to characterise reactivity of matrin-3 (MH3) homology domain antibody, C-terminal antibody, and extreme C-terminus antibody. Related mouse C275 and C181 protein fragments are also shown. Below, dot blot of synthetic peptides, plus bacterially expressed murine fragment C275, showing antibody reactivity. C) Summary of location of epitopes of CIZ1 antibodies in relation to CIZ1 domains. D) Epitope availability of CIZ1 in Xi SMACs in murine (3T3), and female human (HMEC, MCF7) cells, comparing five anti-CIZ1 antibodies by immunofluorescence. Right, histogram shows frequency of CIZ1 SMAC detection in cycling populations.

**Supplemental Figure 5.**
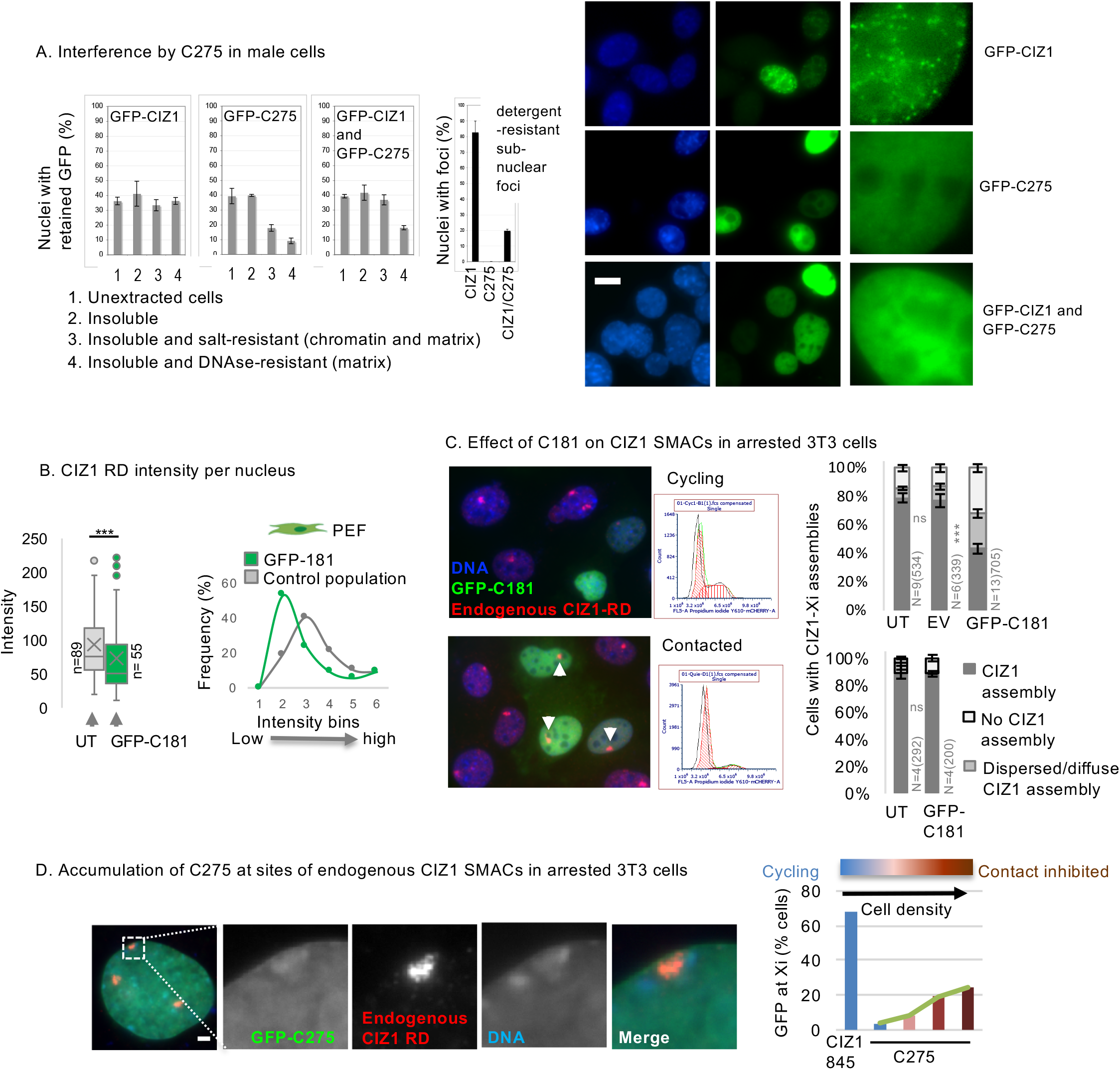
Extended cell cycle analysis. A) Effect of C275 on GFP-CIZ1 sub-nuclear assembly formation in male NIH3T3 cells. Left, graphs show the proportion of cells with nuclear fluorescence (± SEM) after transfection with GFP-CIZ1, GFP-C275 or equivalent amounts of each vector as a co-transfection for 24 hours, with and without exposure to a series of solubilizing buffers prior to fixation (Wilson et al., 2016). GFP-C275, but not full length-CIZ1, is sensitive to extraction of chromatin with DNAse1, and in co-transfection interferes with GFP-CIZ1 from forming DNAse1-resistant assemblies. Black histogram and images (right) show inhibition of assembly of GFP-CIZ1 into foci, by C275 (N=3). GFP fluorescence (green) after extraction with 0.05% Triton X-100 is shown. DNA is blue, bar is 10µm. B) Endogenous CIZ1 fluorescence intensity per nucleus in WT female PEFs. Left, box and whisker plot showing mean, median and interquartile range in untransfected (UT) and C181 transfected cells, where n is nuclei measured. Mann-Whitney U test p=0.00087. Right, frequency distribution of intensity values showing a fall in endogenous CIZ1 intensity in cells expressing C181 (green). C) Effect of C181 in cycling and contact inhibited female D3T3 cells. Left, field images showing both untransfected and transfected cells, illustrating effect on endogenous CIZ1 status at Xi (red). Arrows point to resistant CIZ1 Xi assemblies in contact inhibited cells. Middle, flow cytometry profiles of populations stained with propidium iodide, illustrating G1/G0 enrichment in the contacted cell population. Right, SMAC frequency. For cycling cells (upper graph), no difference in frequencies were observed between untransfected (UT) and empty vector (EV) cells (for CIZ1 marked Xi category p=0.72, dispersed/diffuse p=0.19, no CIZ1 SMAC p=0.75), but dispersal was observed in C181 expressing cells compared to UT (for CIZ1 marked Xi category p=1.4×10^-5^, dispersed/diffuse p=0.0036, no CIZ1 SMAC p=0.00032). For contacted cells (lower graph), none or limited differences in frequencies were observed between UT and C181 expressing cells (CIZ1 marked Xi category p=0.098, dispersed/diffuse CIZ1 patch p=0.34, no CIZ1 marked Xi p=0.043). N=replicate analyses, n=nuclei scored. Error bars show SEM. All comparisons of replicate analyses are by unpaired t-test. D) Frequency of cells with C275 enrichment at sites of endogenous CIZ1 SMACs in female D3T3 cells in serial dilution cultures, showing increase with cell density and contact inhibition (N=1). Example image shows C275 transfected cell, where DNA is blue, GFP green, and endogenous CIZ1 red. Bar is 1µm.

**Supplemental Figure 6.**
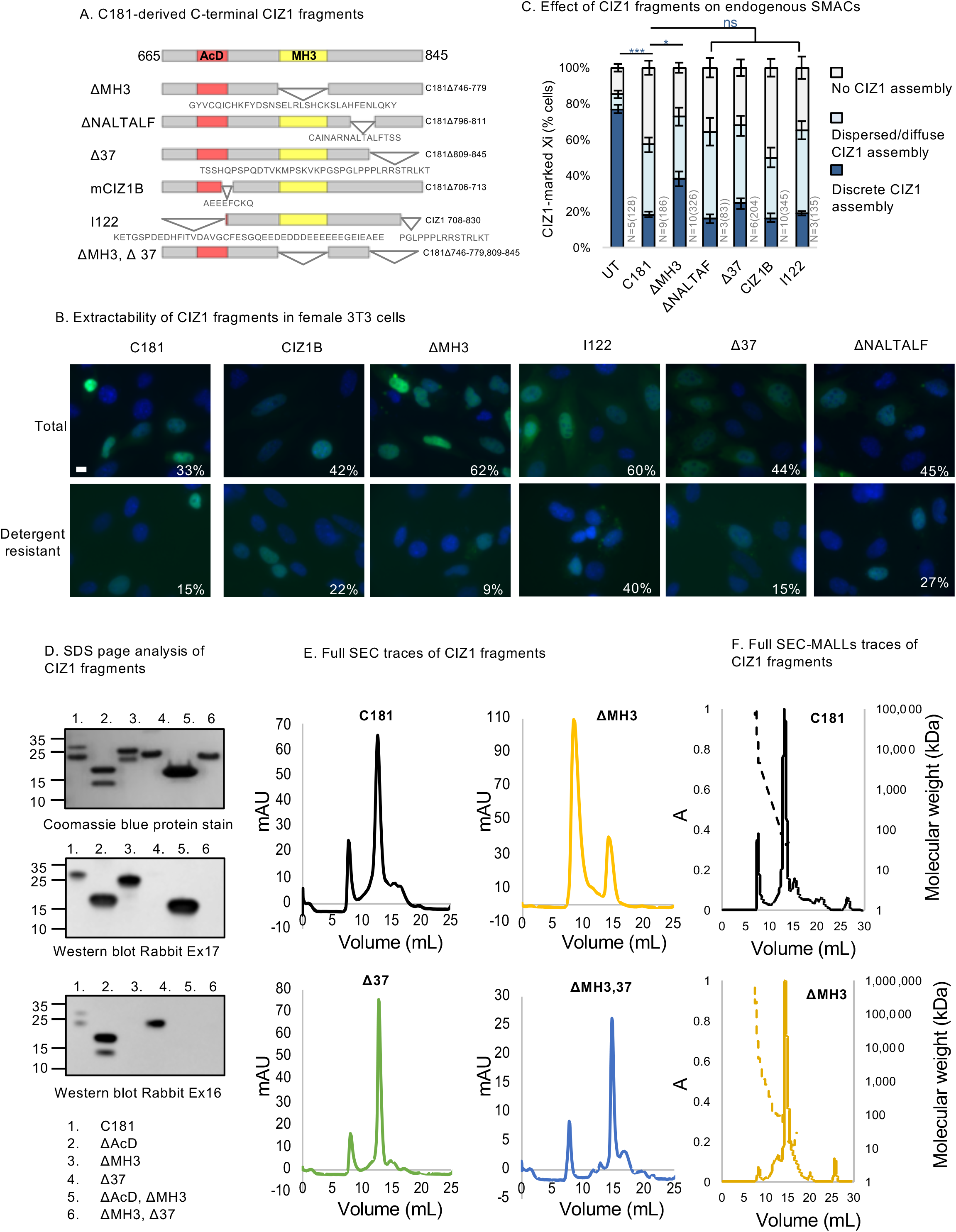
CIZ1 mutant characterisation. A) Map of C181 deletion constructs, showing excluded sequences in single letter amino-acid code. Numbers indicate amino-acid at boundaries relative to murine full-length CIZ1. AcD, acidic domain (red), MH3, matrin 3 homology domain (yellow). B) Field views of female D3T3 cells expressing GFP-tagged C181-derived deletion mutants, with (detergent-resistant) and without (total) pre-fixation treatment with detergent (0.05% Triton X-100 in CSK). The percentages show nuclear localization and the proportion of green cells in each population, revealing the degree of sensitivity to extraction. Bar is 5μm. C) Effect of fragments on the frequency of endogenous CIZ1 SMACs in female D3T3 cells. Compared to C181, only ΔMH3 was perturbed in its ability to disperse endogenous CIZ1 assemblies. For UT to C181 p=9.2×10^-7^, for C181 to ΔMH3 p=0.011. All other deletion mutants retained similar dispersal capability to C181 (ΔNALTALF p=0.96, Δ37 p=0.64, CIZ1B p=0.99, I122 p=1). N shows replicate analyses with total nuclei inspected in parentheses. Comparisons are by one-way ANOVA with a Games-Howell post hoc test. Error bars show SEM. D) Purified recombinant C181, and derived mutant proteins, resolved by denaturing SDS-PAGE. Upper, coomassie blue stain of total protein. C181 shows typically retarded migration (relative mobility at 28/23 kDa compared to the expected encoded 20kDa), as does ΔMH3 (relative mobility at 26/21kDa, compared to the expected encoded 16 kDa), while deletion of the ΔAc domain reverts to the expected position in the gel (relative mobility at 18/14kDa compared to the expected encoded 18kDa). C181, ΔAcD and ΔMH3 all present with a higher and a lower molecular weight band associated with its full-length form, and a cleaved version. The most C-terminal 37 amino-acids is vulnerable and partially cleaved during expression, yielding mixed preparations. Experimental deletion of the 37 most C-terminal amino acids results in a single (retarded) species. Middle, western blot with rabbit Ex17 antibody showing lack of reactivity with cleaved forms and with Δ37 forms. Lower, western blot with rabbit Ex16 antibody with epitope in the MH3 domain, which is reactive with both full-length product and cleaved forms, but not with the ΔMH3 mutants. Antibody epitopes are shown in SFig.4. E) Individual complete size exclusion chromatography (SEC) traces for C181 and C181 mutants, as shown in Fig.4A. All traces have an earlier peak corresponding to the void volume as well as a later peak corresponding to the CIZ1 entity. F) Individual complete size exclusion chromatography multiple angle laser light scattering (SEC-MALLs) traces of C181 and ΔMH3, as shown in Fig.4B. Solid lines are the UV_280_ measurements to indicate protein presence, and the dotted lines are light scattering measurements used to determine the molecular weight of the entities.

**Supplemental Figure 7.**
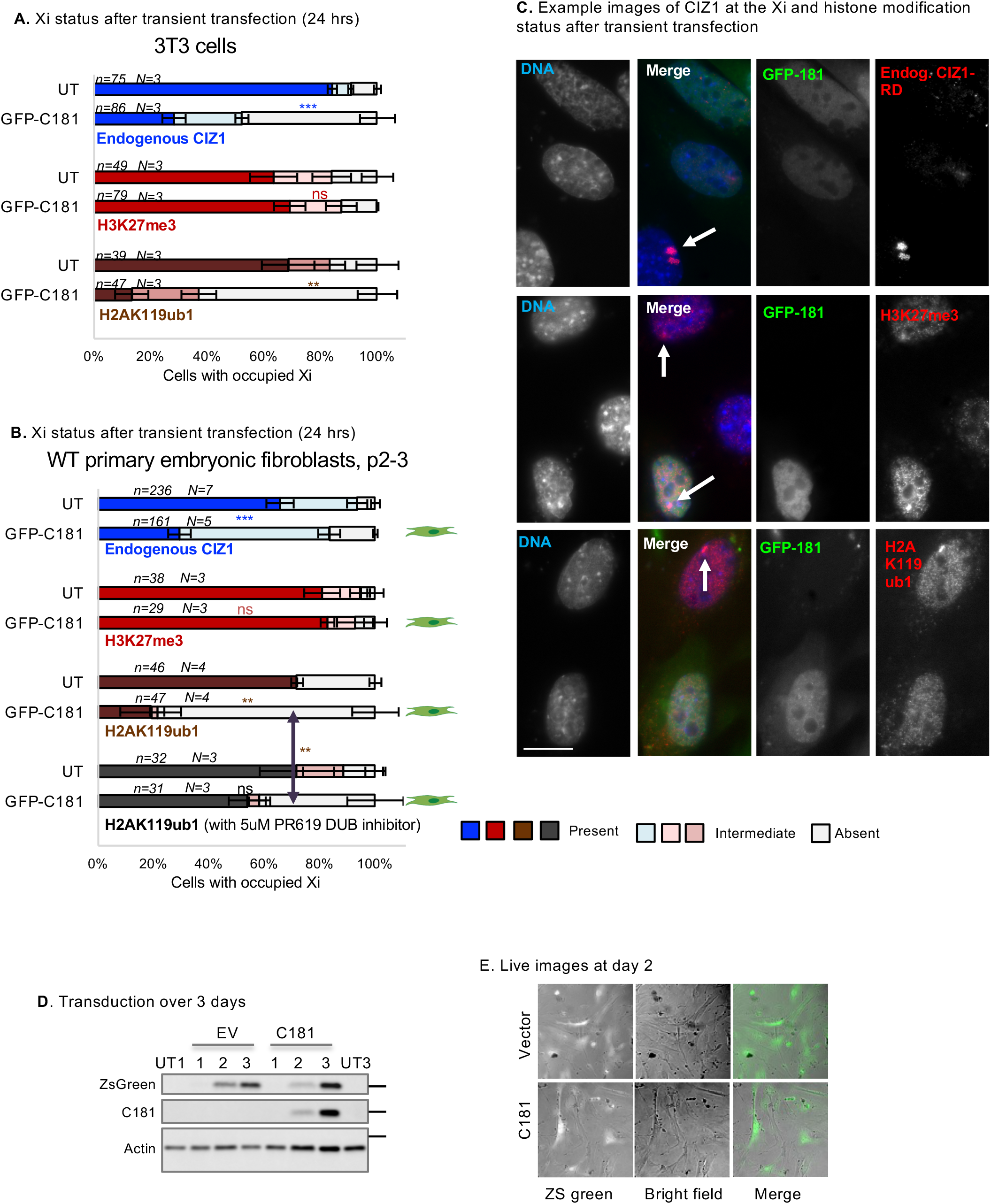
Effect of C181 on CIZ1, H3K27me3 and H2AK119ub1. A) Graphs show the frequency of endogenous CIZ1 SMACs in a cycling population of female D3T3 cells, comparing transfected and untransfected cells in the same population. Endogenous SMACs are detected via CIZ1-RD and classified into three categories; present, absent or intermediate. Middle and lower graphs show the frequency of repressive histone marks in cells that are, or are not, transfected with GFP-C181. N is replicate analyses, n is nuclei scored. Comparisons are by t-test. For endogenous CIZ1 in UT and C181 cells p=0.00023, for H3K27me3 p=0.60, for H2AK119ub1 p=0.0073. Error bars show SEM. B) As in A, except that all data is derived from analysis of female primary embryonic fibroblasts (PEFs) at passage 2-3. For endogenous CIZ1 in UT and C181 cells p=0.00033, for H3K27me3 p=0.79, for H2AK119ub1 p=0.016, performed on present categories. Also shown, the effect of 5μM PR619 on H2AK119ub1 loss, where p=0.0099 for the no CIZ1 category. Error bars show SEM. C) Example images of endogenous CIZ1 and histone marks (red) in untransfected and transfected (green) WT PEF populations. Bar is 10μm. D) Western blot of whole cell lysates from WT female PEFs (13.27p2) 1-3 days post transduction with empty vector control (EV) or with C181, plus untreated control populations (UT) at days 1 and 3. Detection of ZsGreen confirms transduction and expression. Ectopic CIZ1 is detected with rabbit exon 17 antibody only in the C181 transduced group. Beta-actin is loading control. E) Brightfield images and ZsGreen fluorescence in live PEFs at day 2 post transduction.

**Supplemental Figure 8.**
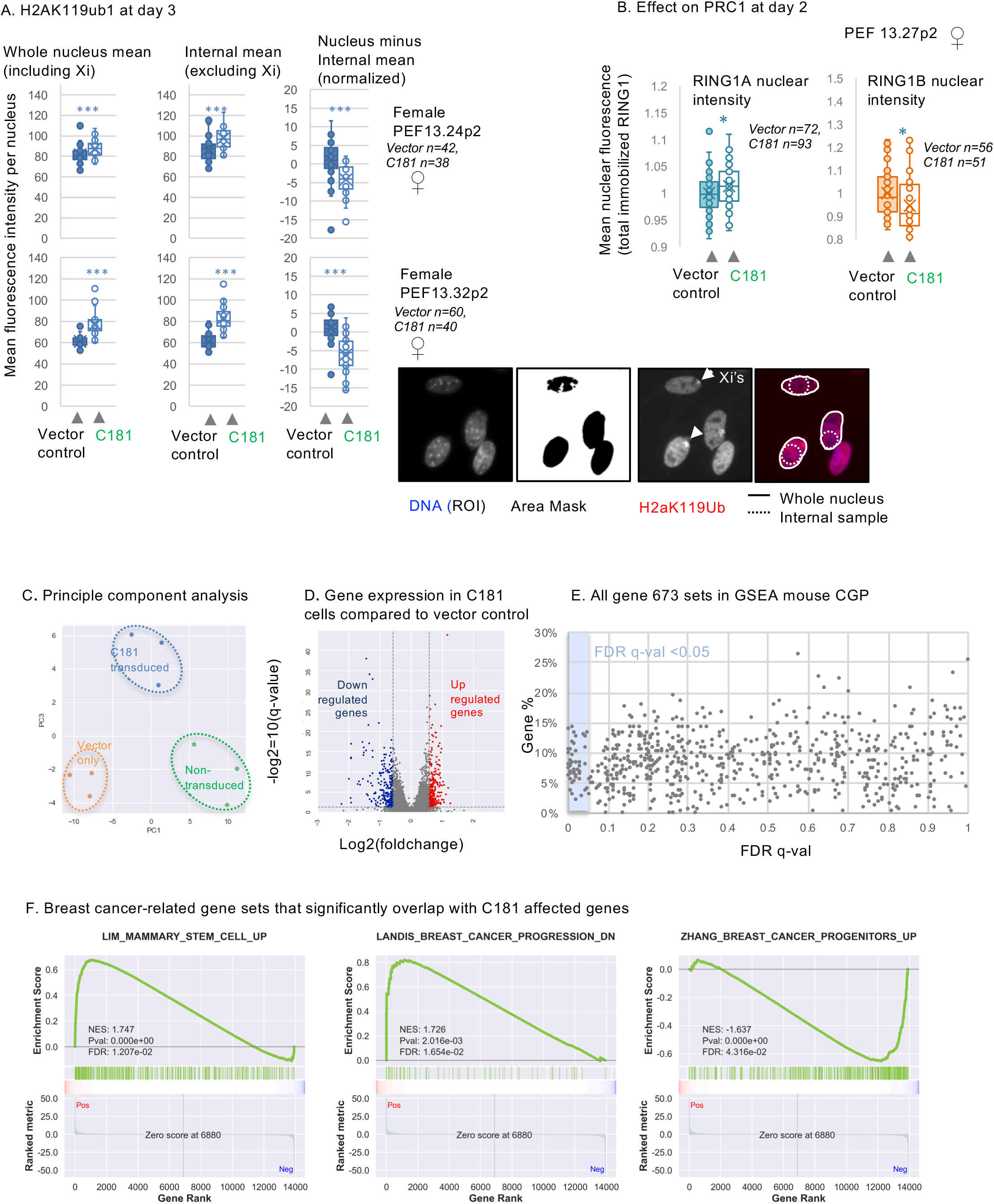
Analysis of transduced cells. A) Analysis of H2AK119ub1 nuclear fluorescence intensity in two female WT PEF populations at day 3 post-transduction. Mean values for the whole nucleus (with inclusion of Xi, left), or a sample area excluding Xi (middle), and subtracted values for each nucleus (right) are shown. Method is illustrated to right. For both populations the data show that C181 drives H2AK119ub1 fluorescence up across the nucleus, but proportionally less at the Xi. B) Mean fluorescence intensity per nucleus of detergent-resistant RING1A and RING1B in female PEFs (N=1), showing C181-induced changes. C) Principle component analysis showing separation of triplicate samples for each condition. D) Volcano plot showing transcription units that are significantly affected by C181. E) Gene set enrichment analysis performed on pre-ranked transcription units comparing expression in C181 transduced cells to the empty vector control, showing all gene sets in the GSEA molecular signature chemical and genetic perturbations. q values are plotted against % genes in overlap, and those with q of <0.05 in the shaded section to left (also in Fig.5G). Source data is given in Supplemental data set 2. F) Example GSEA output showing significant enrichment between three breast cancer-related gene sets and C181 affected genes.

**Supplemental Table 1.**
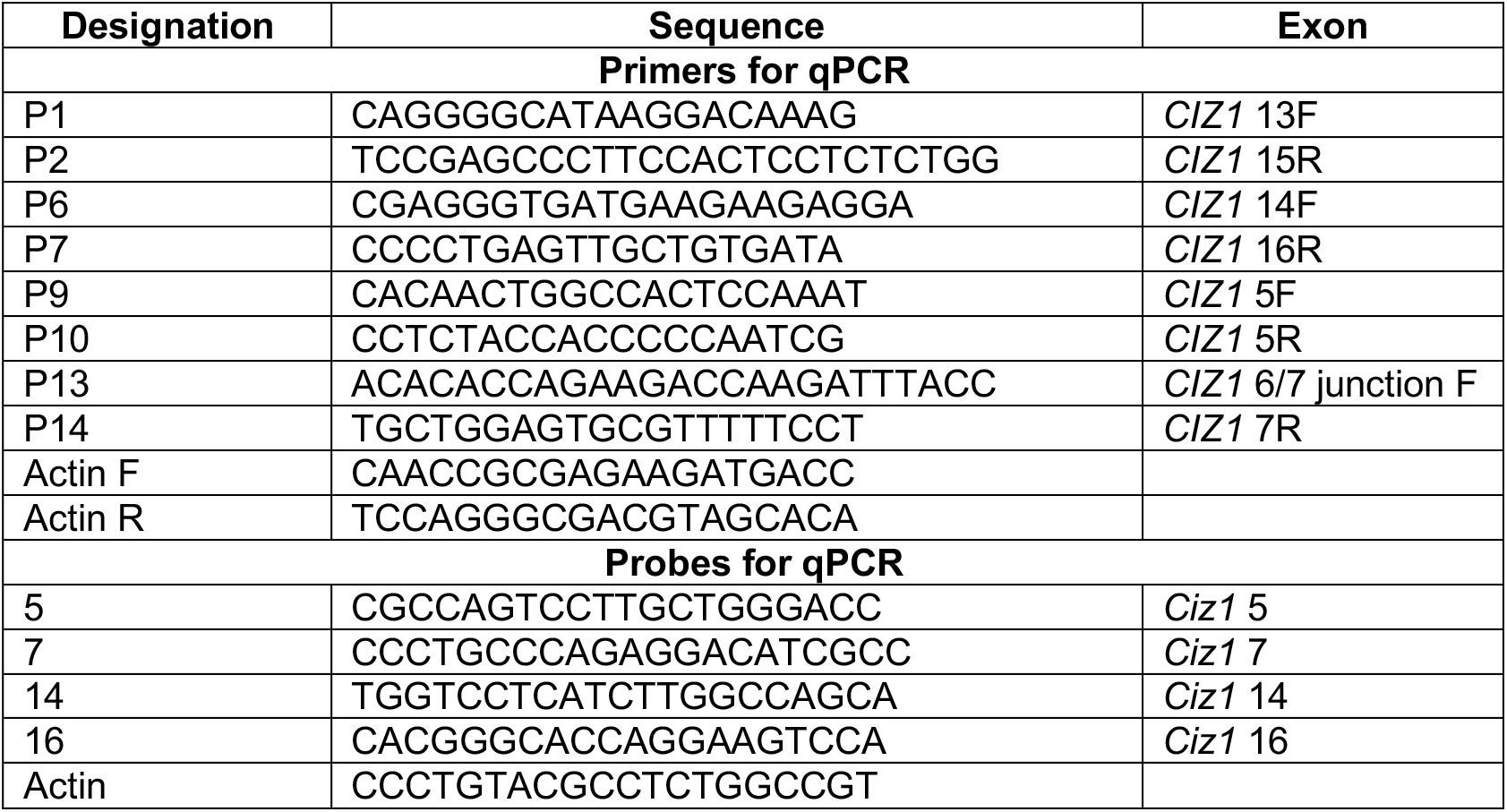
Primer and probe sequences. Primers 9 and 10 were combined with a probe in exon 5 to generate detection tool set DT5, primers 13 and 14 with probe 7 (DT7), primers 1 and 2 with probe 14 (DT14), and primers 6 and 7 with probe 16 (DT16).

**Supplemental Table 2.** Cell line authentication evidence. (Eurofins Medigenomix Forensik GmbH)

|  | **Cell line** |  |  |  |  |
| --- | --- | --- | --- | --- | --- |
| **Marker** | **BT-474** | **MDA-MB-231** | **MCF-7** | **MCF 10A** | **SKBR** |
| AMEL | X,X | X,X | X,X | X,X | X,X |
| CSF1PO | 10,11 | 12,13 | 10,10 | 10,12 | 12,12 |
| D2S1338 | 19,19 | 20,21 | 21,23 | 21,26 | 20,25 |
| D3S1358 | 17,17 | 16,16 | 16,16 | 14,18 | 17,17 |
| D5S818 | 11,13 | 12,12 | 12,12 | 10,13 | 9,12 |
| D7S820 | 9,12 | 8,8 | 8,9 | 10,11 | 9,12 |
| D8S1179 | 10,12 | 13,13 | 10,14 | 14,16 | 11,12 |
| D13S317 | 11,11 | 13,13 | 11,11 | 8,9 | 11,12 |
| D16S539 | 9,11 | 12,12 | 11,12 | 11,12 | 9,9 |
| D18S51 | 13,18 | 11,16 | 14,14 | 18,19 | 10,13 |
| D19S433 | 14,17 | 11,14 | 13,14 | 13,15 | 14,14 |
| D21S11 | 28,32.2 | 30,33.2 | 30,31 | 28,30 | 30,30.2 |
| FGA | 22,25 | 22,23 | 23,24,25 | 22,24 | 20,20 |
| TH01 | 7,7 | 7,9.3 | 6,6 | 8,9.3 | 8,9 |
| TPOX | 8,8 | 9,9 | 9,12 | 9,11 | 8,11 |
| vWA | 15,16 | 15,16 | 14,15 | 15,17 | 17,17 |

**Supplemental Table 3.** Antibodies.

| **Antibodies** | **Working Dilution** | **Source** |
| --- | --- | --- |
| **Immunofluorescence** |  |  |
| N-term ClZ1 (1793/4) | 1 in 1,000 | (Coverley et al., 2005) |
| C-term ClZ1 (87) | 1 in 20 | FDAB |
| C-term ClZ1 (79) | 1 in 50 | FDAB |
| ClZ1 Rabbit Ex16 | 1 in 2000 | Biorbyt (329770) |
| ClZ1 Rabbit Ex17 | 1 in 1,000 | Novus (NB100-74624) |
| Rabbit H3K27me3 | 1 in 1,000 | CST (9733S) |
| Rabbit H2AK119ub1 | 1 in 250,000 | CST (8240) |
| Goat $\alpha$ Rabbit Alexa Fluor 568 (Red) | 1 in 1,000 | Invitrogen (A11011) |
| Goat $\alpha$ Mouse Alexa Fluor 488 (Green) | 1 in 1,000 | Invitrogen (A11001) |
| Goat $\alpha$ Rabbit Alexa Fluor 488 (Green) | 1 in 1,000 | Invitrogen (A11034) |
| Goat $\alpha$ Mouse Alexa Fluor 568 (Red) | 1 in 1,000 | Invitrogen (A11031) |
| **Western Blot** |  |  |
| N-term ClZ1 (1793/4) | 1 in 1,000 | (Coverley et al., 2005) |
| ClZ1 Rabbit Ex16 | 1 in 200 | Biorbyt (329770) |
| ClZ1 Rabbit Ex17 | 1 in 1,000 | Novus (NB100-74624) |
| H3 | 1 in 5,000 | Abcam (ab1791) |
| β-Actin | 1 in 2000 | Abcam (ab11003) |
| Peroxidase IgG $\alpha$ Rabbit | 1 in 10,000 | Jackson 211-032-171 |
| Peroxidase IgG $\alpha$ Mouse | 1 in 10,000 | Jackson 155-035-174 |

**Supplemental Table 4.**
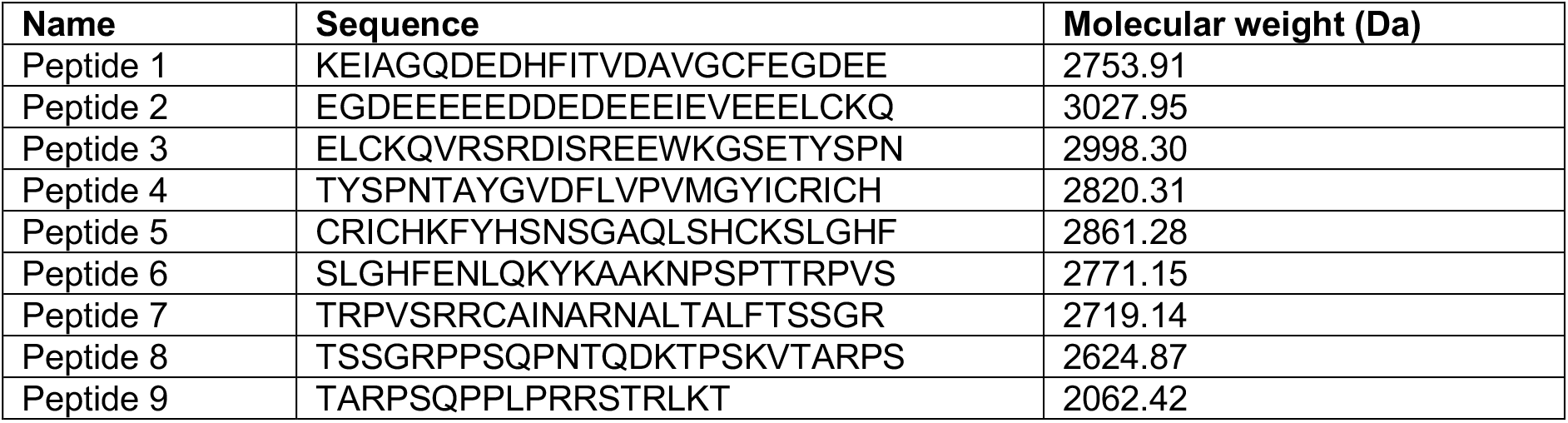
Peptides.

**Supplemental Table 5.**
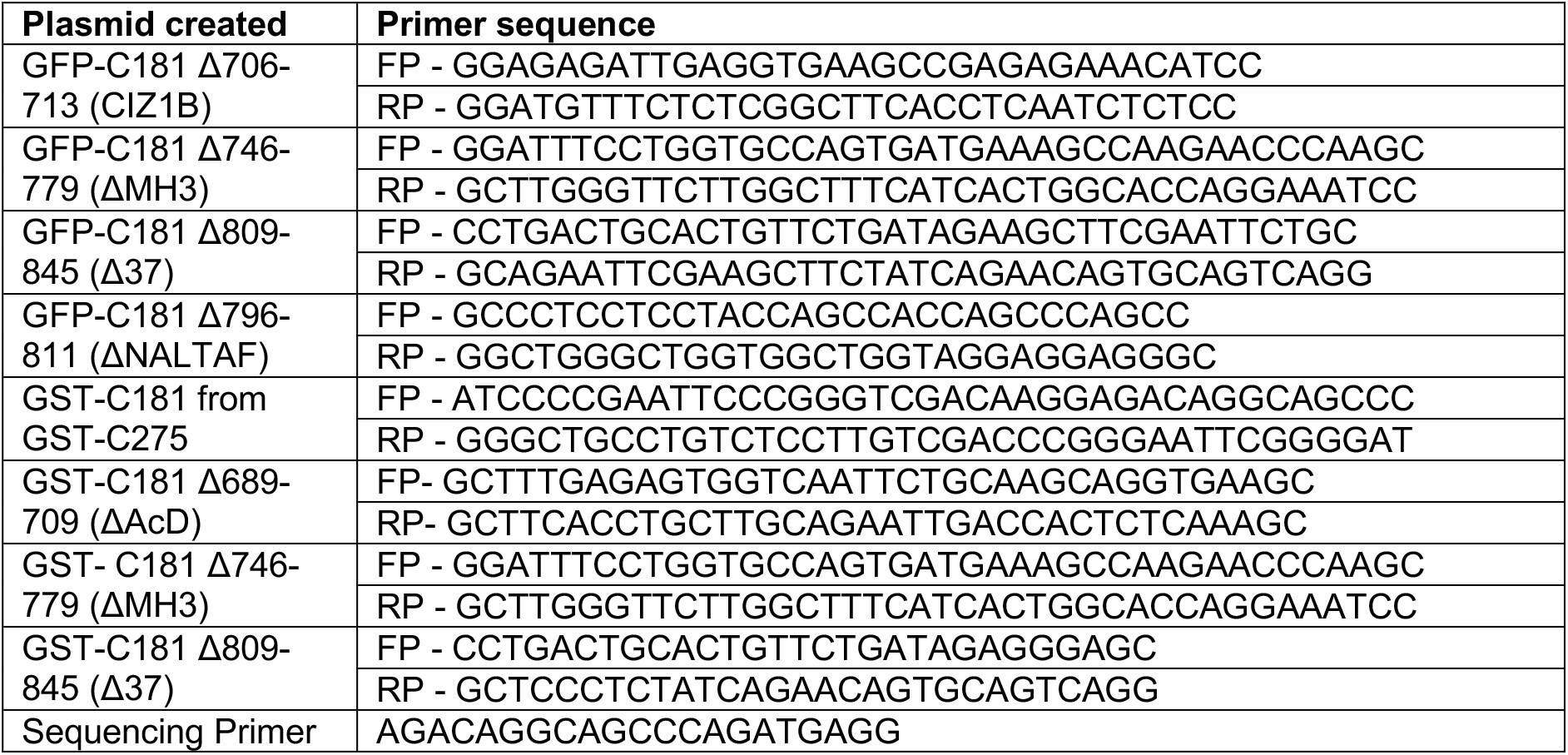
Primers used in mutagenesis.

**Supplemental data set 1 (Xl file)** Raw data and primary tumour designations for QRTPCR analysis.

**Supplemental data set 2 (Xl file)** Transcriptomics and GSEA analysis

